# snoRNA-guided tRNA 2’-O-methylation controls codon-biased gene expression and cellular states

**DOI:** 10.1101/2023.06.22.546166

**Authors:** Minjie Zhang, Kongpan Li, Jianhui Bai, Ryan Van Damme, Wei Zhang, Mario Alba, Bangyan L. Stiles, Jian-Fu Chen, Zhipeng Lu

**Author notes:** Author emails: M.Z., K.L., J.B., R.V.D, M.A., B.L.S., J.-F.C., Z.L.

## Abstract

snoRNAs are a large family of noncoding (nc)RNAs present across eukaryotes and archaea. While a subset of them guide 2’-O-methylation (Nm) and pseudouridylation (Ψ) of rRNAs and snRNAs, targets of most snoRNAs remain unknown. Here we used PARIS2 to map snoRNA targets, revealing an extensive and conserved snoRNA-tRNA interaction network. Using optimized denatured RiboMeth-seq (dRMS), we discovered snoRNA-guided Nm sites in ncRNAs, including tRNAs. Loss of snoRNAs and their associated 2’-O-methyltransferase FBL reduced tRNA modifications and increased fragmentation. CRISPR knockout of the D97/D133 family of snoRNAs reduced the activity and levels of several target tRNAs, including elongator (e)Met-CAU, leading to codon-biased transcriptome and translatome in human cells. The codon-biased gene expression tipped the balance between the dichotomous cellular states of proliferation and differentiation, and skewed germ layer potential of mouse embryonic stem cells. Together, we discovered a snoRNA-guided tRNA modification mechanism controlling codon-biased gene expression and cellular states.

## INTRODUCTION

Small nucleolar (sno)RNAs are a large family of noncoding (nc)RNAs present in both eukaryotes and archaea, which split more than 4 billion years ago ^1–3^. The human genome encodes ∼2000 snoRNA genes, many of which are differentially expressed in cell types and developmental stages ^4, 5^. Most snoRNAs are classified into two types based on the presence of conserved sequence and structure features. snoRNAs often use antisense guide sequences upstream of D/D’ or H/ACA motifs to recognize targets ^6–11^. C/D snoRNAs bind four proteins, including the 2’-O-methyltransferase (MTase) FBL, to catalyze 2’-O-methylation (Nm). H/ACA snoRNAs bind four other proteins, including the pseudouridine synthase DKC1, to catalyze pseudouridylation (Ψ). While some well-studied snoRNAs localize to the nucleolus to guide rRNA modifications, a small fraction that reside in the Cajal body, termed small Cajal (sca)RNAs, target spliceosomal snRNAs. Archaeal homologs are called small (s)RNAs due to lack of the nucleolus. All are commonly referred to as snoRNAs for convenience. Beyond the rRNAs and snRNAs, snoRNAs have been shown to bind tRNAs ^3, 12–17^, snoRNAs, other types of ncRNAs ^18^, and likely mRNAs ^19–25^. The vast majority of snoRNAs have no identified targets and are often referred to as orphans. Even for snoRNAs with known targets, their initial discoveries were based on biased predictions and may be incomplete. Therefore, the targets of all snoRNAs await comprehensive discovery and validation.

In recent years, genetic analysis in metazoa revealed critical roles for snoRNAs and their associated enzymes in physiology and pathology. For example, FBL levels regulate *C. elegans* lifespan and host-pathogen interactions ^26–28^. DKC1 mutations cause a genetic disorder dyskeratosis congenita and predisposes to cancer ^29, 30^. Genetic screens identified several snoRNAs that control cholesterol metabolism, lipo/gluco-toxicity, and oxidative stress ^31–36^. Loss of *SNORD116*, an imprinted cluster, causes Prader-Willi syndrome ^37^. Mutations in *SNORD118/U8* cause leukoencephalopathy, a neurological disorder ^38–42^. Increasing evidence demonstrated critical roles of snoRNPs in tumorigenesis ^43–51^. However, our limited knowledge of snoRNA targets made it difficult to study their molecular mechanisms in these biological contexts.

In addition to carrying amino acids for translation, tRNAs also act as sequence-specific regulators of cellular states^52^. The tRNA pool is dynamically adjusted in development and in response to the environment via tissue-specific expression, chemical modification, splicing and charging (aminoacylation). Together, these mechanisms produce specific tRNA pools that control various processes, such as mRNA stability, codon-biased translation, and altered protein folding kinetics leading to differential functional or defective products ^53, 54^. Upon stress, tRNAs can be cleaved into small fragments (tRFs) with translation-independent functions, such as stress response, extracellular signaling, and transposon silencing ^55–57^. Dysregulation of tRNA biogenesis factors causes a variety of cancers and genetic disorders ^58^. tRNAs are among the most densely modified RNAs, with more than 100 chemical varieties, and up to 1 out of 6 nts is modified in eukaryotes. These modifications are essential for function and are disrupted in numerous human diseases, yet the modification mechanisms remain poorly understood, despite decades of research ^59–61^.

In this study, we use PARIS2, a highly optimized method for in vivo RNA duplex detection ^41^, to discover a large number of new snoRNA target sites on rRNAs, snRNAs, several other ncRNAs, as well as nearly all nuclear-encoded or cytoplasmic (cyto)-tRNAs, but not mitochondrial (mt)-tRNAs. The C/D snoRNAs bind pre-tRNAs, suggesting that their guided modifications occur early in tRNA biogenesis. While previous studies have found a single eukaryotic snoRNA-guided tRNA Nm site in eMet tRNA ^15^, our analysis revealed a much larger and deeply conserved network. Developing an optimized denatured RiboMeth-seq (dRMS) method, we validated and discovered snoRNA-guided Nm sites in ncRNAs, including tRNAs. Re-analysis of published CLIP data showed that multiple C/D and H/ACA snoRNP proteins bind most human and yeast cyto-tRNAs. Loss of specific snoRNAs, such as the SNORD97/D133 (D97/D133) family, and the MTase FBL, reduced Nm and other modifications, and increased tRNA fragmentation. CRISPR knockout (KO) of the D97/D133 family reprogrammed the transcriptome and translatome to adapt to the reduced activity and level of D97/D133 target tRNAs, including the eMet-CAU. This codon-biased gene expression dictates the balance between the dichotomous cellular states of proliferation vs. differentiation/development/morphogenesis. In a mouse embryonic stem cell (mES) differentiation model, the D97/D133-dependent codon-biased gene expression controls proliferation and germ layer specification, especially the differentiation to cardiomyocytes (CM). Together, this study revealed a snoRNA-guided tRNA modification mechanism that controls codon-biased gene expression and cellular states.

## RESULTS

### PARIS2 and dRMS capture new snoRNA target sites across multiple ncRNAs

C/D and H/ACA snoRNAs base pair with targets to guide chemical modifications (**Fig. 1A-B**). To discover snoRNA targets, we designed biotinylated antisense oligos (bio-ASOs) to enrich 46 snoRNA families, including 36 orphans (see STAR Methods). We used psoralens to crosslink RNA base-pairing interactions in human cell lines HEK293, SH-SY5Y, and induced pluripotent stem cell (iPS)-derived lineages, such as astrocytes, endothelial cells (EC), neuro-progenitors (NPC), and neurons ^40, 41^. PARIS2 was used to discover snoRNA targets in total RNA, bio-ASO-enriched snoRNA complexes and chromatin-associated RNAs (caRNA) ^41^ (**Fig. 1C**). To validate newly discovered targets, we used published RMS data ^62, 63^, developed an optimized denatured RMS (dRMS, **Fig. S1A- J**), and analyzed published snoRNP CLIP data for target enrichment and chimeric reads that support interactions ^18, 64^ (**Fig. 1D**).

**Figure 1.**
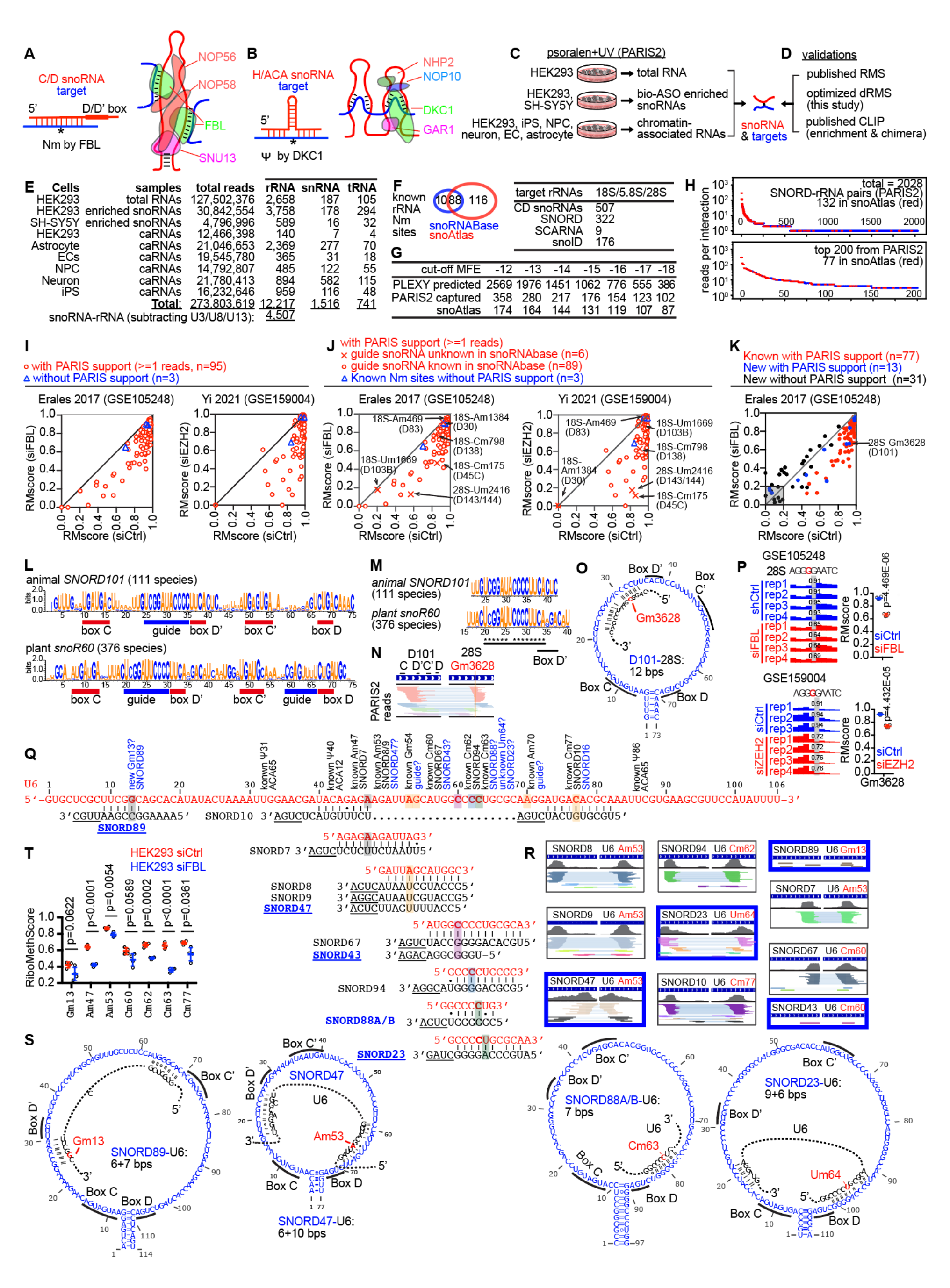
PARIS2 and dRMS discover new snoRNA targets. (A-B) Models of C/D and H/ACA box snoRNP complexes. (C-D) Strategies for the PARIS2 identification of snoRNA targets and validation. (E) PARIS2 captured three major types of snoRNA targets. At the bottom, snoRNA-rRNA interactions were summed after removing interactions with 3 snoRNAs U3/U8/U13, which guide rRNA processing, but not chemical modifications. (F) Known Nm sites on rRNAs summarized from two databases (left) and guide snoRNAs (right). Total known and predicted rRNA-targeting C/D snoRNAs (507) include 322 SNORD, 9 SCARNA, and 176 newly annotated (snoID) in snoAtlas. (G) Numbers of SNORD-rRNA interactions from PLEXY predictions, PARIS2, and snoAtlas at various MFE (kcal/mol) cutoffs. (H) Ranked SNORD-rRNA duplex groups (DGs) captured by PARIS2. Red dots are snoAtlas annotations. (I) RMscores for 98 known rRNA Nm sites, based on published data. PARIS2 missed 18S-Cm463, 28S-Um4499 and 28S-Gm4500. (J) Data are the same as panel I. PARIS2 identified guide snoRNAs for all 6 known Nm sites, for which guide snoRNAs were previously unknown in snoRNABase (in parentheses, e.g., D83 is short for SNORD83). Among the 6, D83 was predicted to guide 18S-Am468 ^75^, D103 was predicted to guide 18S Gm601 ^76^, D45C was recently confirmed to modify Cm175 ^77^. (K) Discovery of new rRNA Nm sites from Erales 2017 RMS ^62^. Nm sites are defined as p<0.05 in RMscore differences between siCtrl and siFBL (n=121), regardless of whether the changes were positive or negative. Out of them, 77 are supported by PARIS2 data (>=1 reads). Out of the 121, 13 were previously unknown, but supported by PARIS2, out of which, 10 have decreased RMscores after FBL KD, while 3 have increased RMscores. Out of the 121, 31 were previously unknown and not supported by PARIS2, out of which, 12 have decreased RMscores after FBL KD, while 19 have increased RMscores. (L-M) Alignments of chordate and plant D101 homologs (L) and D’ guide sequences (M), based on Rfam clan CL00074. (N-O) PARIS2 chimeric reads (N, 90 reads) and structure model (O) supporting the 28S-D101 interaction. (P) RMS data supporting the predicted Gm3628 and its loss upon FBL and EZH2 KD. P values: unpaired two-sided t-tests. (Q) Human U6 snRNA sequence (red) and PARIS2 support for known and new snoRNA-snRNA interactions. Blue text: either new modifications or newly discovered interactions that mediate known modifications. Guide snoRNAs for several known Nm sites, including Gm54 and Am70, are still missing. In addition to its known target of 18S rRNA, the SNORD16 interaction with U6, potentially responsible for the Cm77 modification, was discovered in our previous work ^78^. Am53 is targeted by at least 3 different snoRNAs, D8 (known), D9 (known) and D47 (newly discovered). (R) Example snoRNA-U6 interactions supported by PARIS2. New interactions are highlighted in blue boxes. (S) Example structure models for new interactions and Nm sites. Cm63 is not supported by PARIS2 reads, but supported by dRMS. Three of the examples are bipartite interactions. (T) dRMS validated Nm sites on U6 snRNA. Um64 is not modified based on dRMS and excluded. See also Figure S1 and Tables S1-S3.

PARIS2 revealed multiple types of snoRNA targets, including rRNAs, snRNAs, tRNAs, 7SL, and snoRNAs themselves (**Fig. 1E**, **Table S1**). C/D snoRNA targets were captured more efficiently than H/ACA snoRNA targets. Here, we focused on C/D snoRNAs for initial validation. PARIS2 captured significant fractions of potential targets among PLEXY-predicted low energy interactions and known targets in published databases at each minimal free energy (MFE) cutoff (**Fig. 1F-G, Fig. S1K**). Among 2028 PARIS2- captured snoRNA-rRNA interactions with modification guide snoRNAs (excluding U3/U8/U13), 132 are consistent with the snoAtlas database ^4^ (**Fig. 1H**). The rest of them are likely new functional interactions, or transient contacts that represent the scanning of snoRNPs on targets ^65^. In the top 200 PARIS2-captured interactions, 77 are consistent with the snoAtlas database. Out of 98 known rRNA Nm sites in snoRNABase, the vast majority have reduced Nm levels upon disruption of snoRNPs by FBL and EZH2 knockdown (KD) ^62, 63^, and PARIS2 captured 95 of them (**Fig. 1I**). For known Nm sites in rRNAs, guide snoRNAs were either unknown or only predicted in the snoAtlas database (n=6), which were all discovered by PARIS2 (**Fig. 1J, Table S2**). De novo discovery of potential sites where Nm levels are reduced after FBL KD also revealed previously unknown sites, a subset of which are supported by PARIS2 (blue, n=13, **Fig. 1K**). For example, D101 is highly conserved in animals and plants, especially in the D’ guide (**Fig. 1L-M**). PARIS2 and structure modeling revealed a strong interaction targeting 28S Gm3628 (**Fig. 1N-O**). Analysis of published RMS data confirmed the reduction of Nm level after FBL or EZH2 KD (**Fig. 1P**) ^62, 63^.

The commonly used RMS hydrolyzes RNA at high pH, where 2’-O-methylation protects the phosphodiester bond, which is detected by sequencing ^66–68^. However, stable RNA structures and dense modifications cause strong biases in the fragmentation and reverse transcription, impeding its application to many ncRNAs. We developed a denatured RMS (dRMS) method by using alkaline pH, high temperature and DMSO, which increased the efficiency and uniformity of RNA fragmentation (see STAR Methods, **Fig. S1A-J**). Combining PARIS2 and dRMS, we validated known and discovered new Nm sites on several small RNAs, including spliceosomal snRNAs, 7SL, and snoRNAs (**Fig. 1Q-T, Fig. S1L-Z, Table S3**). For example, U6 is a heavily modified snRNA ^69^. Guide snoRNAs were only known for a subset of them (**Fig. 1Q**). PARIS2 captured guide snoRNAs for known Nm sites, and potential new Nm sites (blue boxes) (**Fig. 1R-T**). Among them, D89 and D23 were orphans, D47 also targets the 28S rRNA, whereas D43 also targets the 18S ^4^. These studies confirmed the accuracy of PARIS2 and revealed new snoRNA targets across multiple ncRNA types.

### PARIS2 and dRMS reveal a global snoRNA-tRNA interaction network

In addition to rRNAs and snRNAs, cyto-tRNAs are a major group of snoRNA targets (**Fig. 2A, Fig. S2A,** and **Table S4**). While some snoRNAs bind only rRNAs or tRNAs, others can bind both, suggesting functional pleiotropy (**Fig. 1N-P**, **Fig. S2B-D, Table S1**). For C/D snoRNA-target chimeric reads, fragments mapped to the snoRNAs are piled around the expected D/D’ guide regions (**Fig. 2B- C**). To determine whether PARIS2-captured interactions are energetically favorable, we shifted the tRNA-mapping fragments to each side of the identified target sites (**Fig. 2D**). The PARIS2 chimeras, but not randomly shuffled ones, produced a deep MFE valley at the target sites (**Fig. 2E-F**). Together, these two analyses support the validity of the snoRNA-tRNA network. Furthermore, tRNA fragments in snoRNA-tRNA chimeras often extend beyond the mature tRNA transcript. suggesting that the interactions occur on pre-tRNAs, prior to removal of the leader and trailer sequences (**Fig. 2G**).

**Figure 2.**
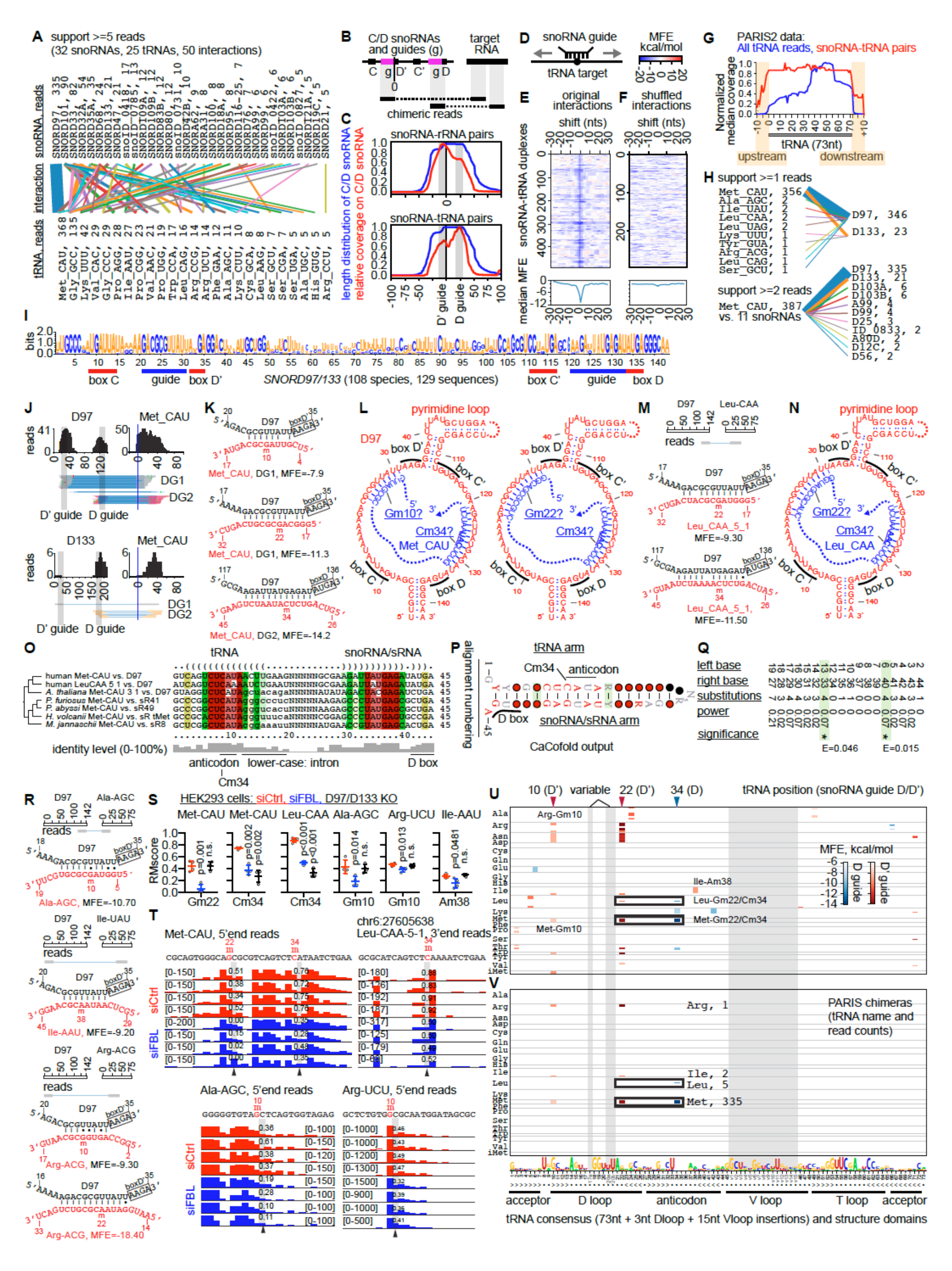
PARIS2 and dRMS reveal a global network of snoRNA-tRNA interactions. (A) The global network of snoRNA-tRNA interactions supported by >=5 reads and ranked by chimeric support (PARIS2 + CLIP). Line thickness is scaled to the square root of read numbers, which are listed after the RNA name. (B) For all chimeras connecting C/D snoRNAs, the coverage was averaged over the snoRNAs. The example chimeras have one arm (left side) mapped to the D’ and D guide motifs. For the meta-gene analysis, the position after the D’ guide is set to 0. (C) For all snoRNA-target pairs, coverage of the arms mapped to snoRNAs are summed up in red. Blue lines are the average length distribution of snoRNAs (x axis: nucleotides). Positions for the average 10nt D’ and D guides are labeled. (D) For all C/D snoRNA-tRNA duplexes detected by PARIS2 and CLIP, each duplex was shifted to the left or right of the target Nm site for 30 single-nucleotide steps, and MFE was calculated for each step. (E) PARIS2 DGs were shifted for MFE calculation (top, n= 490). MFE medians were calculated for each position (bottom). (F) Same as panel E, except that randomly positioned snoRNA-tRNA interactions on the experimentally determined snoRNA-tRNA pairs (n=297 successful shuffles) were used to generate the matrix (top) and medians (bottom). (G) Coverage of all PARIS2 reads mapped to tRNAs were normalized to max=1, including continuous reads (non-gapped), intra-molecular structure reads, and intermolecular interaction reads. For all snoRNA-tRNA interactions, arms mapped to tRNAs were summarized in red, normalized to max=1, revealing high coverage up-and down-stream of the standard 73nt tRNA gene model. (H) D97/D133 and eMet-CAU snoRNA-tRNA sub-networks. (I) D97/D133 homologs from Rfam were aligned using MAFFT, retaining positions in >=30% sequences, and visualized in WebLogo. (J) PARIS2 reads supporting D97/D133 interactions with eMet-CAU tRNA. Blue vertical lines represent the start of the mature tRNA. D’ and D guide sequences are highlighted in gray boxes. (K-L) Models of D97 interactions with eMet-CAU (MFE in kcal/mol). DG1 supports two alternative conformations with the D’ guide. (M-N) PARIS2 gapped reads and models of D97 interactions with Leu-CAA-5-1 tRNA (MFE in kcal/mol). (O) LocARNA structured alignments of eMet-CAU and Leu-CAA-5-1 interactions with guide RNAs. Human eMet-CAU and Leu-CAA-5-1 are intron-less. The A. thaliana eMet-CAU-3-1 and archaeal eMet-CAU tRNAs have introns, in which the first 8 nts were shown. The other 8 intron-less A. thaliana eMet-CAU tRNAs, not shown here, cannot base pair with the D97 homolog due to sequence variations. Only two types of A. thaliana tRNAs contain introns, including 11 eMet-CAU genes, and 61 Tyr-GUA genes. (P-Q) R-scape and CaCofold analysis of the interactions between eMet-CAU / Leu-CAA tRNAs and their guide RNAs across eukaryotes and archaea. Alignments are numbered 1-45 (20nt tRNA, 5nt N and 20nt guide). Panel Q shows the statistics for all base pairs proposed by R-scape. Gray boxes and asterisks indicate significant covariation above evolutionary background. (R) PARIS2 gapped reads for additional D97 snoRNA targets, secondary structure models and MFE (kcal/mol). (S-T) dRMS analysis of D97/D133 target tRNAs in siCtrl, siFBL and D97/D133 double KO HEK293 cells. RMscore: riboMeth-seq score. P values: unpaired two-sided t-tests. n.s.: not significant. n=4 samples for each group. Example dRMS read end pileup tracks are illustrated for siCtrl and siFBL samples (T). The 5’ and 3’ end counts were used, whichever has higher coverage for that tRNA. (U-V) Heatmap for PLEXY predicted (U) and PARIS2/CLIP discovered (V) Nm sites guided by D97. Each row is a tRNA gene (n=430 rows), grouped by anti-codon. Shades of blue represent D guide targets, whereas shades of red represent D’ guide targets. All tRNAs are aligned to a standard model of 73nts plus insertions at the D-loop (not to be confused with the snoRNA D box/guide) and V-loop. Three most targeted sites in the tRNAs are labeled: 10, 22 and 34, and targeted by D’ and D guides, respectively. See also Figures S2-S3 and Tables S4-S6.

D97/D133, eMet-CAU tRNA, and their partners, form the strongest snoRNA-tRNA sub-network (**Fig. 2H**). CRSSANT ^42^ clustering resolved two DGs connecting the D’/D guides to two distinct regions on eMet-CAU (**Fig. 2J, Fig. S2E**). These DGs form strong duplexes, including alternative ones for the D’ guide interaction, that suggest new Nm sites at Gm10 or Gm22, in addition to the previously reported Cm34 (**Fig. 2K-L**) ^15^. PARIS2 also revealed Leu-CAA-5-1 as a new target for D97/D133, likely due to its homology to eMet-CAU (**Fig. 2M-N, Fig. S2F**). Interestingly, the wobble position N34 in many other tRNAs is methylated by FTSJ1, a stand-alone 2’-O-methyltransferase ^70^. Earlier studies in archaea reported sRNAs that guide tRNA modifications ^3, 71^. Alignments of tRNAs and guides in human, plant *A. thaliana*, and 4 archaeal species revealed a conserved duplex (**Fig. 2O**). Interestingly, the target eMet-CAU tRNAs in plants and archaea have introns in the anticodon loop and participate in the extended duplex, further supporting that pre-tRNAs are snoRNA targets (**Fig. 2G**). R-scape and CaCofold ^72, 73^ revealed two significantly covaried base pairs, in addition to 4 invariable base pairs, confirming deep functional homology among archaeal and eukaryotic guide RNAs for eMet Cm34 (**Fig. 2P-Q**). This analysis also revealed a conserved function for the poorly studied tRNA introns in guiding tRNA modifications ^12, 74^. Other D97-tRNA interactions are also supported by strong duplexes, despite the fewer chimera support (**Fig. 2R**).

The Nm levels at 6 sites in 5 tRNA targets are reduced either in FBL KD or D97/D133 double KO, or both, confirming the extended interaction network (**Fig. 2S-T**). The variable Nm levels and changes upon snoRNP reduction may be due to differences in interaction strength, and/or redundancy of additional guide snoRNAs (**Fig. 2S**). Given the low PARIS2 coverage for some validated tRNA targets, we used computational prediction to find additional targets missed by PARIS2 (**Table S5**). Predicted D97 targets on all 432 human cyto-tRNA genes were aligned to a standard tRNA (**Fig. 2U**), revealing many potential sites in several consistent positions (10, 22 and 34), a subset of which were captured by PARIS2 (**Fig. 2V**). Prediction of D133 targets revealed primarily interactions with the D guide due to divergence of sequence of the D’ guide (**Fig. S2H-I**). In addition to D97/D133, several other snoRNAs also target the eMet-CAU tRNA (**Fig. 2H, Fig. S2J, Table S1**). Together, these data revealed an extensive D97/D133- tRNA sub-network.

We further characterized other snoRNA-tRNA interactions and confirmed their stability by structure modeling (**Fig. S2K-W**). D32A, D33, D34 and D35A, 4 rRNA-targeting snoRNAs encoded in the introns of *RPL13A*, regulate stress response ^19, 20^. PARIS2 and structure modeling revealed tRNAs as another major group of targets (**Fig. S2K-R**). Despite the lower crosslinking efficiency, we discovered a few interactions between H/ACA snoRNAs and tRNAs that may guide Ψ, including the highly conserved TΨC motif (**Fig. S2T-W**). To discover Nm sites in tRNAs de novo, we applied dRMS to control and siFBL HEK293 cells (**Table S3**). Reduced Nm levels upon FBL KD were observed in 149 sites, among which, PARIS2 captured guide snoRNAs for 15 sites (**Fig. S3A-C**). Furthermore, 3 of them are also supported by previously published RMS data, despite their lower quality ^62, 63^. KD of *EZH2*, a known oncogene, reduced tRNA modifications, suggesting a connection of snoRNA-tRNA interactions to cancer. Mutations of yeast genes in snoRNA processing and recycling also led to reduced Nm levels in several tRNAs (**Fig. S3D-J, Table S6**), further supporting the conserved snoRNA-tRNA interactions.

### CLIP confirms snoRNP binding to human and yeast cyto-tRNAs

To determine whether snoRNP proteins also bind tRNAs, we analyzed published 365nm UV PAR-CLIP and 254nm UV eCLIP data for human C/D snoRNP proteins NOP56, NOP58, FBL, and the H/ACA snoRNP protein DKC1 ^18, 79, 80^ (**Fig. 3A-B**). Earlier studies failed to find enrichment of tRNAs due to the lack of proper normalization. To test RNA enrichment in CLIP, we calculated the ratio of read counts between CLIP and a small RNA-seq dataset (20-200nt range). Given the spatial segregation of nuclear snoRNPs and mt-tRNAs, we reasoned that mt-tRNAs can serve as a normalization standard, providing a conservative estimate of the enrichment ratios (**Fig. S4A-E**, STAR Methods). To identify a cutoff for significantly enriched RNAs, we set an empirical false positive rate (FP) at 0.05 for miRNAs, which are not known to interact with any snoRNP (**Fig. 3A**).

**Figure 3.**
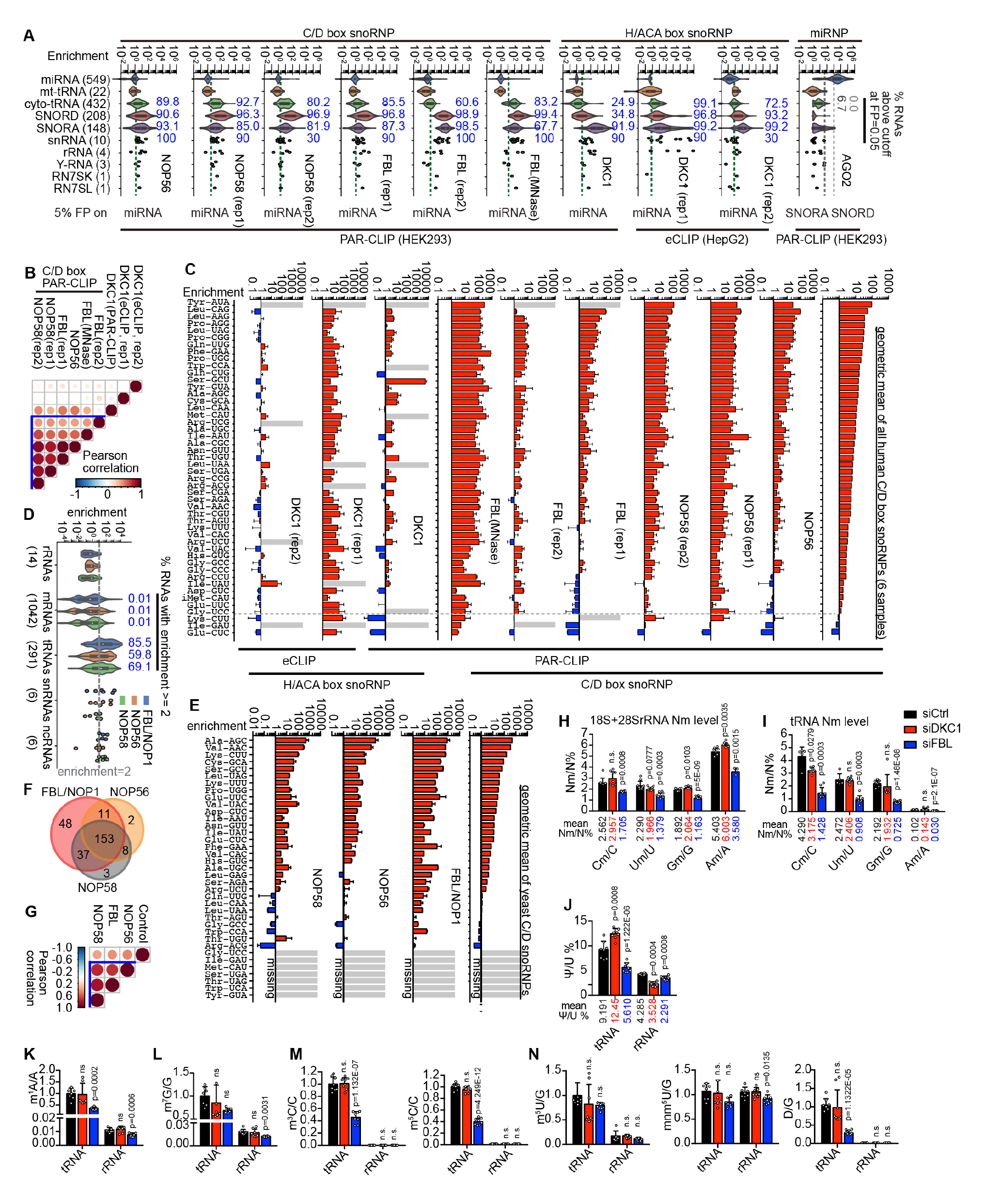
snoRNP complexes bind tRNAs and catalyze their modifications. (A) RNA enrichment ratios in PAR-CLIP over 20-200nt sRNA-seq input, and eCLIP over size-matched input, normalized to the median of 21 mt-tRNAs. Numbers in parentheses indicate genes in each type. Data were plotted in as violins (when n>10) or individual dots. In the box plots in the violins, whiskers represent the max and min. The top and bottom of the box represent the 1st and 3rd quartiles. The circles in the middle represent the medians. miRNAs served as negative controls for snoRNP CLIP where vertical dash lines indicate the ratio above which 5% miRNAs are considered enriched (FP=0.05). For the AGO2 CLIP, the FP=0.05 was defined using SNORD or SNORA RNAs (lighter and darker vertical dash lines). The blue-colored numbers are the % of RNAs above the FP=0.05 cutoff. SNORD, box C/D snoRNAs. SNORA, box H/ACA snoRNAs. (B) Pairwise Pearson correlations among human snoRNP CLIP experiments. (C) Enrichment ratios for each tRNA anticodon group in snoRNP CLIP. Error bars: s.d. for all tRNA genes in each group. Gray bars: missing data. tRNAs were ranked by the geometric means across the 6 CLIPs for NOP56, NOP58 and FBL. (D) Enrichment ratio of yeast RNA relative to input control. Dash line: the 2-fold cutoff for calculating % of enriched RNAs. (E) Enrichment ratios for each anticodon group of yeast tRNAs. Error bars: s.d. across tRNA copies within each isotype. Gray bars: missing data. tRNAs were ranked by the geometric means across the 3 CLIPs. (F) Venn diagram of enriched tRNAs from the 3 yeast CLIP experiments, for tRNAs with RPM>1 and enrichment ratio >=2. (G) Pairwise Pearson correlations among all yeast snoRNP CLIP experiments. (H-J). Quantitative LC/MS analysis of Nm and Ψ levels from isolated total tRNA and 18S/28S rRNAs after DKC1 and FBL KD. P values are from two-sided unpaired t-tests between the siRNA KD and the siCtrl cells. n.s.: not significant. Levels are quantified and adjusted by standard curves. (K) LC/MS analysis of other modifications in total tRNAs and 18S/28S rRNAs. The levels are relative due to the lack of standard curve normalization (ratios set to 1 in siCtrl). See also Figure S4 and Tables S4

Using this approach, we found that C/D snoRNP core proteins enriched C/D and H/ACA snoRNAs, snRNAs and rRNAs, as expected ^18^ (**Fig. 3A**). 7SK and 7SL are also enriched, consistent with previous studies ^18^. Surprisingly, between 60% and 93% of cyto-tRNAs are also enriched, much higher than the empirical 5% FP for miRNAs, and the negative control mt-tRNAs (**Fig. 3B**). H/ACA snoRNP protein DKC1 enriched, C/D and H/ACA snoRNAs, snRNAs and rRNAs, although with lower efficiency ^18^ (**Fig. S4B- C**). Similarly, a subset of cyto-tRNAs were also enriched (25-99%). As another control, AGO2 PAR-CLIP enriched miRNAs, but not any other RNA types (**Fig. 3A**, here FP defined on snoRNAs). Different C/D snoRNP proteins enriched tRNA anticodon groups in similar patterns (**Fig. 3C**). CLIP experiments occasionally produce hybrid reads that indicate spatial proximity of two RNA species ^81, 82^. Re-analysis of CLIP data using CRSSANT revealed a few snoRNA-tRNA chimeras, most of which are consistent with PARIS2 data (**Table S4**).

We further analyzed published 254nm UV CLIP data for yeast C/D snoRNP proteins FBL/NOP1, NOP56 and NOP58 (**Fig. 3D**) ^64^. We calculated enrichment ratios against CLIP data on a non-tagged control yeast strain (Control), and normalized the ratios using reads per million (RPM) after removing the snoRNAs, the major targets of snoRNP proteins (**Fig. 3D**). Likewise, this approach provides a conservative estimate on the enrichment ratios. Setting enrichment ratio = 2 as a cutoff, between 59 and 85% yeast tRNAs were enriched by this standard (**Fig. 2D**), and they are highly consistent across the experiments (**Fig. 3E-G**). Together, both yeast and human CLIP data revealed most cyto-tRNAs as snoRNP targets.

### FBL regulates multiple types of tRNA modifications

Both *FBL* and *DKC1* are essential for cell survival ^83–85^. We performed mass spectrometry on purified tRNA and 18S/28S rRNAs after FBL and DKC1 KD (**Fig. S1I**, **Fig. S4H**). Nm levels were reduced in both 18S+28S rRNAs and tRNAs upon FBL KD, while DKC1 KD did not change Nm levels except Cm in tRNA (**Fig. 3H-I**). Interestingly, we observed larger reductions in Nm levels in tRNAs than in rRNAs after FBL KD. Ψ level was reduced in rRNAs but not tRNAs after DKC1 KD, suggesting either interactions that do not guide tRNA modifications, or only few tRNA Ψ sites are catalyzed by DKC1 (**Fig. 3J**). Surprisingly Ψ was reduced after FBL KD in tRNAs, even though FBL does not have the Ψ synthase activity, suggesting complex regulation of RNA modifications by snoRNPs. Further quantification showed that multiple other tRNA modifications were also reduced in tRNAs upon FBL KD, but not DKC1 KD (**Fig. 3K-N**). Together, these results suggest that C/D snoRNP catalyze Nm modification of tRNAs and then further regulate other tRNA modifications, consistent with the snoRNP binding to pre-tRNAs.

### snoRNA-guided tRNA modifications protect tRNAs from angiogenin cleavage

Earlier studies have identified several modifications that stabilize tRNAs, including the D97/D133-dependent Cm34 in eMet tRNA ^15^. To determine whether snoRNPs protect tRNA in general, we knocked down FBL and DKC1 in two cell lines HEK293 and A549 (**Fig. S1I, Fig. S5A**, **Fig. 4A-H**). FBL KD significantly increased fragments in the 15-50 nt range, either in the absence or presence of oxidative stress (arsenite, **Fig. 4A-B**). The induced fragments include tRNA halves (∼34 and 40 nts) and shorter ones below 20nts from D and T loop cleavage. In contrast, DKC1 KD did not induce tRFs (**Fig. 4C-D**). To validate these effects and rule out side effects of siRNA transfections, we generated stable HEK293, A549 and HepG2 cell lines expressing shRNAs targeting FBL and DKC1 (**Fig. S5B-G**), and subject the cells to arsenite oxidative stress, alkaline pH 9.0, and heat shock. In all these conditions, shRNA KD of FBL induced more tRFs without apoptosis. To determine whether the increased fragmentation was due to defects on tRNAs per se or secondary changes in cellular physiology, we purified tRNAs from wildtype and siFBL cells, and incubated them with purified endonuclease angiogenin (ANG), which typically cleaves the tRNA anticodon loop (**Fig. 4I-L**). tRNAs from FBL KD cells are significantly more susceptible to cleavage, generating a wide range of sizes, most of which are tRNA halves ^86^. The in vitro melting temperatures of purified total tRNAs were not changed after FBL KD, indicating that overall tRNA structures were intact (**Fig. 4M**).

**Figure 4.**
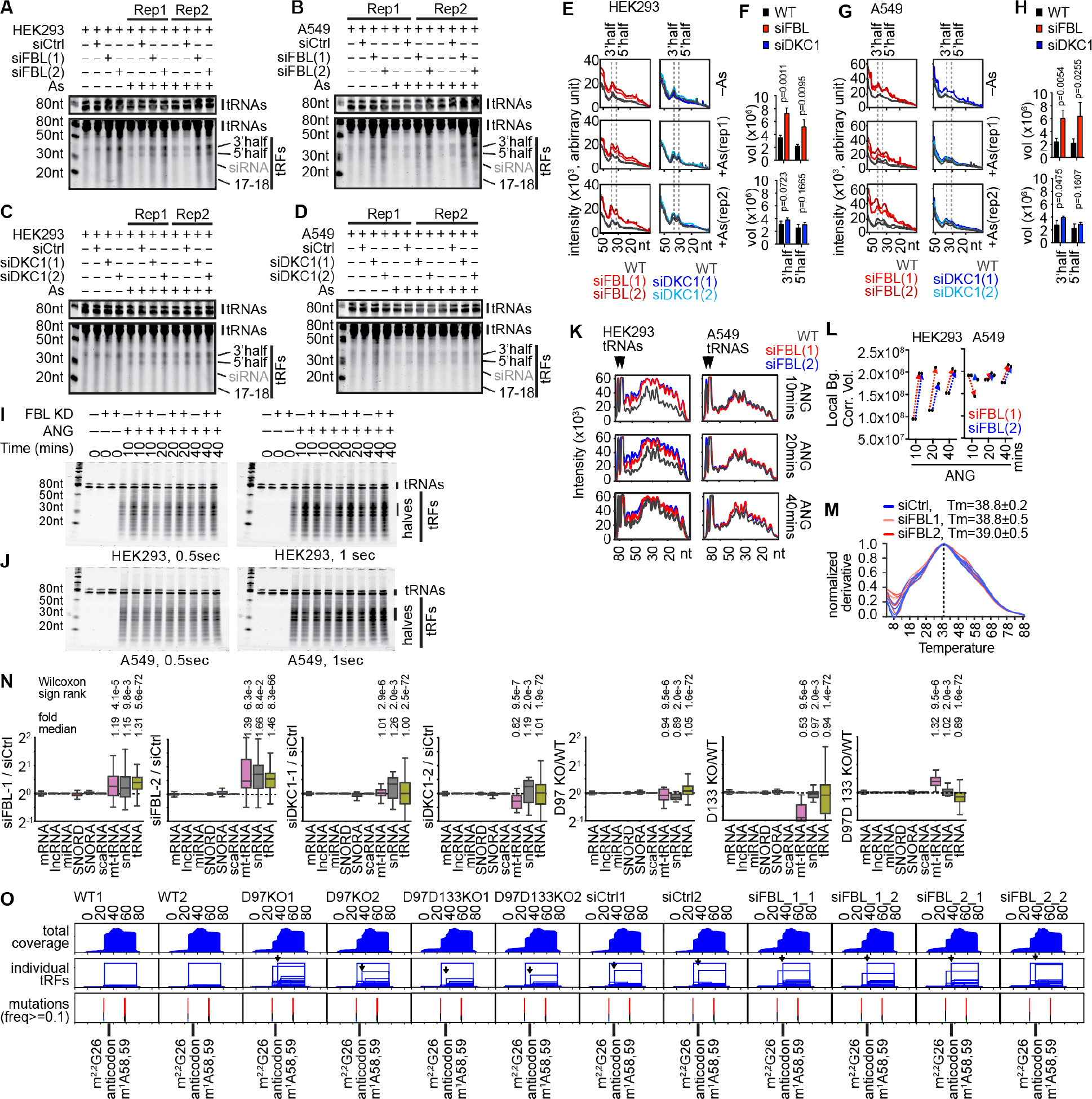
FBL and snoRNAs protect tRNAs against fragmentation. (A-D) SYBR Gold stained total RNA from HEK293 and A549 cells after FBL or DKC1 KD and 4 hours of 0.25mM sodium arsenite (As) treatment. Transfected siRNAs are occasionally visible on the gels. Top panels: shorter exposures of tRNA bands. (E-H) Quantification of RNA fragments from HEK293 and A549 (panels A-D). tRNA halves were quantified for the bars (F, H). Error bars are ± s.d. of n=2 independent siRNA experiments. (I-J) tRNA in vitro cleavage assay. tRNA bands, ∼70-90 nucleotides, from HEK293/A549 cells were isolated from total RNAs after electrophoresis, and digested by recombinant human ANG. (K-L) Quantification of tRNA fragments from HEK293 and A549 cells (panels I-J). (M) Melting temperature of total tRNAs from 10-day siCtrl and siFBL HEK293 cells measured on a thermocycler and SYBR Green. Technical replicates n=5. First derivative was calculated and normalized to max=1. (N) Changes of 15-50nt RNA levels in siFBL and siDKC1 cells. Y-axis: RNA RPM (reads per million) ratios in KD vs. siCtrl, or KO vs. WT. Each value was added 50 to avoid zero divisions and reduce variation, e.g., (RPMsiFBL-1+50) / (RPMsiCtrl+50). (O) Met tRFs from HEK293 cells. Individual tRFs were extracted and plotted as rectangles. Each rectangle is one group of identical reads defined by the start and end, where height is the number of reads. Arrow: start the 3’ tRNA half. Bars in the mutation track are modified positions that induce RT stops and mutations. Mutation signatures are color-coded (A: green, C: blue, G: black and T: red). See also Figure S5.

To identify tRFs, we sequenced RNA in the 15-50nt range from wildtype, siRNA control, KD of FBL/DKC1, and snoRNA KO strains, after AlkB demethylation of RNA ^87, 88^ (**Fig. S5H**). Compared to siRNA control and siDKC1, siFBL increased global levels of cytosolic tRFs (**Fig. 4N, Fig. S5I-J**). Since these data were normalized to RPM, the actual fold changes should be higher. Interestingly, snRNA and mt-tRFs also increased significantly. The increased fragments are mostly below 40 nts (**Fig. S5I**). To determine the effects on eMet-CAU tRNA, we quantified fragments from the KD and KO strains. Both FBL KD and D97/D133 KO increased eMet-CAU tRNA halves and several other tRFs relative to full length tRNAs (**Fig. 4O**, **Fig. S5K-L**). Together, these studies showed that cellular RNAs, especially tRNAs, are protected by snoRNA-guided 2’-O-methylation, confirmed earlier work on the D97/D133 snoRNAs ^15^, and extended the finding to most cyto-tRNAs.

### D97/D133 KO reprogrammed the Met-AUG codon-biased transcriptome and translatome

Both single and double KO of D97/D133 significantly reduced cell growth (**Fig. 5A**). snoRNA overexpression rescued the defects, ruling out the possibility of disrupting host genes and other off-target effects of the CRISPR method (**Fig. 5B**). Labeling of nascent peptides by the alkyne conjugated Met amino acid analog HPG revealed dramatically reduced global translation in all the KO strains (**Fig. 5C-D**). Then we performed RNA-seq and ribosome profiling (ribo-seq) ^89^ in WT and KO cell lines to measure the specific defects in gene expression (**Figs. S6A-D**). Single KO strains did not change tRNA levels, even though each paralog is necessary for tRNA modifications, however, the double KO reduced expression of several tRNAs, including eMet, Leu, Ile, etc. (**Fig. 5E)**, which are targets of D97/D133 (**Fig. 2**). On the other hand, Pro tRNAs, which are not D97/D133 targets, were significantly upregulated, consistent with a previous study of codon usage reprogramming ^54^ (**Fig. S6E**), suggesting secondary effects on the tRNA pool.

**Figure 5.**
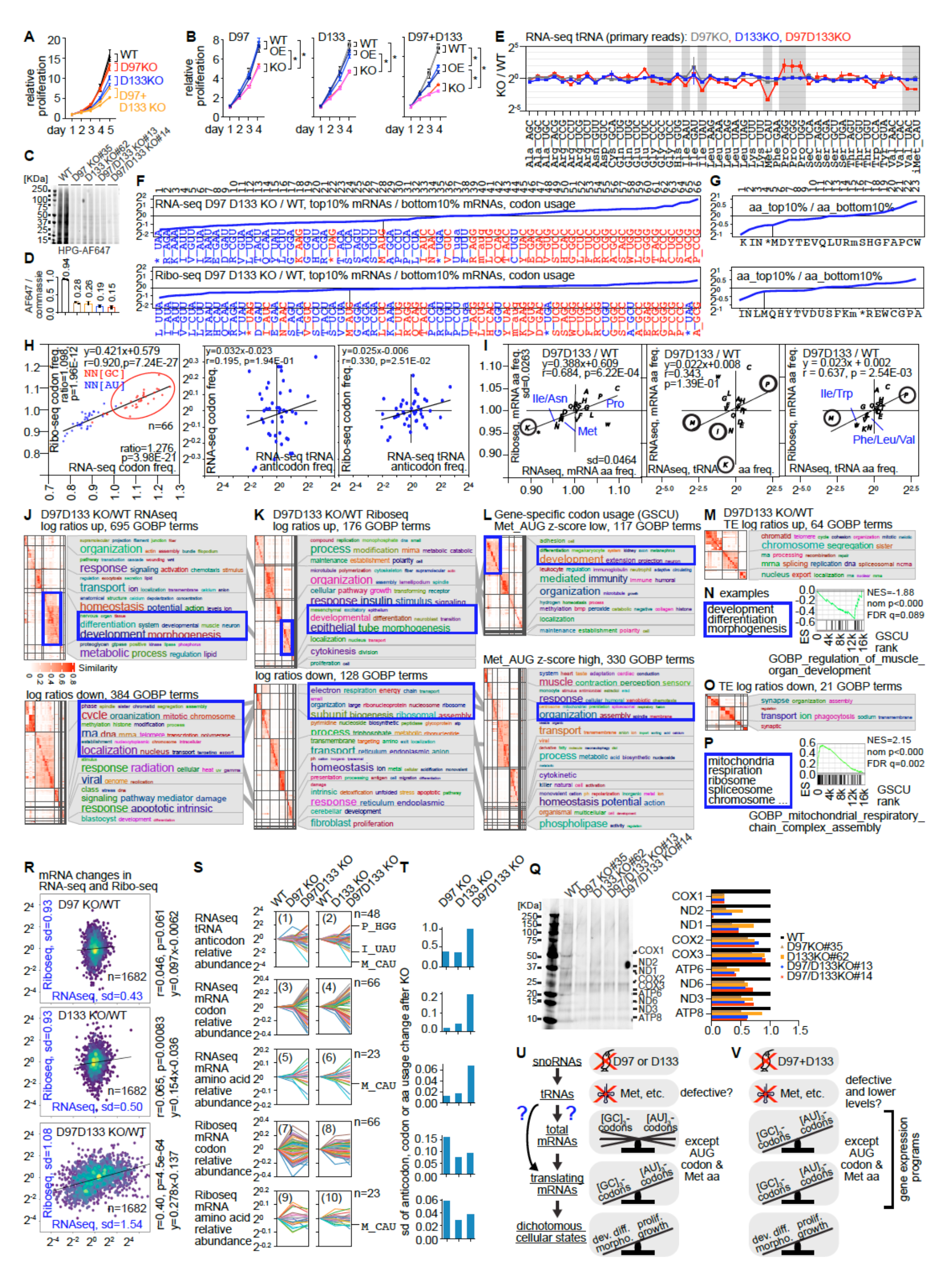
Loss of D97/D133 snoRNAs leads to codon-biased transcriptome and translatome in human HEK293 cells. (A) Relative proliferation of WT and single and double KO HEK293 cells were measured over 5 days (two clones each). P values are listed as follows, based on two-sided unpaired t-tests. P values for D97 KO vs. WT: 0.30 and 0.38 (day 4), 0.0024 and 0.0084 (day 5). P values for D133 KO vs. WT: 0.0005 and 0.0004 (day 4), 3.2E-05 and 3.6E-06 (day 5). P values for D97/D133 double KO vs. WT: 0.0001 and 5.8E-05 (day 4), 2.4E-06 and 2.5E-06 (day 5). (B) KO cell lines, two clones each, were rescued by snoRNA overexpression (OE). P values <0.01 are indicated in the figure (asterisks *, for all clones), based on two-sided unpaired t-tests. (C-D) Nascent proteins were labeled using L-homopropargyl-glycine (HPG), detected using Alexa-Fluor-647 azide (C), quantified and normalized against Coomassie Blue staining of total protein on the gel (D). Two replicates were included for each cell line. (E) All 432 nuclear-encoded tRNA genes were grouped by anticodons and then log2 transformed ratios between KO and WT were normalized so that median=1 for each KO/WT comparison. Error bars are ± s.d. of independent RNA-seq experiments. (F) For nuclear-encoded mRNAs in RNA-seq (upper panel) or ribo-seq (lower panel) from WT and D97/D133 double KO HEK293 cells, ratios of expression levels were ranked. Usage of 66 codons – standard 64 plus iMet and Sec – were calculated for the top 10% (up-regulated) and bottom 10% (down-regulated) mRNAs and weighted by expression levels. Blue and red: A/U- and G/C- ending codons. Asterisk (*): stop codons. M and m: eMet and iMet. U_uga: Sec. (G) Same as panel D, except that amino acid usage was calculated from RNA-seq (upper panel) or ribo-seq (lower panel). n=23 for 20 amino acids plus iMet, stop, and Sec. (H) Changes in codon and anticodon usage were calculated between D97/D133 double KO and WT. RNA-seq codon: usage was calculated for 66 codons in mRNAs from KO and WT and the ratios were normalized to median=1. Ribo-seq codon: same as the RNA-seq codon analysis, except that ribo-seq was used. RNA-seq tRNA anticodon freq: tRNA levels were measured in the RNA-seq as in panel C. Left panel: comparison of RNA-seq and ribo-seq codon usage. NN[GC] and NN[AU]: G/C and A/U ending codons. The inset x-axis ratio=1.276 indicates ratio of average NN[GC] vs. average NN[AU] in RNA-seq. The inset y-axis ratio=1.098 indicates the ratio of average NN[GC] vs. average NN[AU] in ribo-seq. p values after the ratios are from two-sided unpaired t-tests between the two codon groups. Linear regression: fit equation, correlation coefficient r and statistical significance. Middle panel: RNA-seq mRNA codon freq. vs. RNA-seq tRNA codon frequency. Codons recognized by the same tRNA anticodons were merged. Right panel: same as the middle panel, except that ribo-seq mRNA codon frequency is the y-axis. (I) Same as panel H, except that the amino acid usage was plotted. (J) Altered mRNAs based on log ratios of D97D133 KO vs. WT were median-normalized and tested for gene set enrichment using GSEA and MSigDB c5.all collection. Enriched gene sets were clustered based on term similarity and tested for enrichment of terms. GOBP term clusters consistent across RNA-seq, ribo-seq and GSCU were highlighted in blue boxes and linked by gray lines. (K) Same as panel J, except that the analysis was performed on ribo-seq mRNA levels. (L) Same as panel J, except that the analysis was performed on Met-AUG codon enrichment in mRNAs. (M) Same as panel J, except that the analysis was performed on translation efficiency (TE, mRNA levels in ribo-seq over RNA-seq). (N) Example GOBP terms with significantly depleted Met-AUG usage (n=15423 genes with APPRIS principal isoforms). (O) Same as panel J, except that the analysis was performed on TE. (P) Example GOBP terms with significantly enriched Met-AUG usage. (Q) After inhibition of cytosolic translation by cycloheximide, the alkyne-containing methionine homolog HPG was used to label nascent peptides ^92^. Then mitochondria were isolated from cells and labeled proteins were reacted with Azide-Alexa647 and visualized on a gel (left). Mitochondrial peptides were quantified (right). (R) Ratios of mRNA levels in KO vs. WT were plotted for RNA-seq and ribo-seq. Standard deviations (sd) were calculated for RNA-seq and ribo-seq, respectively. Numbers of mRNAs plotted were listed (e.g., n=1682 for D97 KO/WT). Linear regression results on the right: correlation coefficient r, statistical significance p and linear fit equation. (S) Nuclear-encoded tRNA anticodon groups (n=48), mRNA codon (n=66), and mRNA amino acid (n=23) usage changes upon D97/D133 single and double KO. Panels 1-2: Relative abundance of tRNA anticodons in KO vs. WT (WT set to 1). Panels 3-6: mRNA codon and corresponding amino acid usage on the transcriptome level. Panels 7-10: mRNA codon and corresponding amino acid usage on the translatome level (ribo-seq). (T) Standard deviations (sd) of anticodon, codon and amino acid usage plots in panel R. (U-V) Model for the effects of snoRNA-guided modifications on the tRNA pool, mRNA codon usage, translation, and cellular states. Single (U) and double (V) KO affect the transcriptome and translatome to different degrees. See also Figure S6.

In both the transcriptome and translatome (ribosome-associated mRNAs), the usage of many codons, including Met-AUG, changed significantly in the double KO (**Fig. 5D-E**). There is a clear enrichment of GC-ending codons and depletion of AU-ending ones (**Fig. 5E**), which has been correlated with stem cell self-renewal and differentiation (**Fig. S6F**) ^54^, and multicellular functions ^53^. On the amino acid level, Met usage is the fourth most reduced in the double KO in both the transcriptome and translatome (**Fig. 5F-G**).

Strong positive correlations were observed between the transcriptome and translatome in the codon and amino acid usage changes in double KO vs. WT (**Fig. 5H-I**). The D97 and D133 single KO did not show the same trend of codon and amino acid frequency changes on the transcriptome, but changed codon and amino acid frequency on the translatome level, suggesting that the eMet tRNA is also defective in the single KO cell lines (**Fig. S6G-J**).

We calculated up and down-regulated gene ontology (GO) terms ^90^, and clustered them to identify enriched ones in a word cloud ^91^. All KO strains exhibited significant downregulation of genes involved in basic cellular functions, such as ribosome, spliceosome, mRNA metabolism, and oxidative phosphorylation (**Fig. 5J-K, Fig. S6K-M**), consistent with the reduced growth and translation. GO terms upregulated on the transcriptome and translatome levels include differentiation, development, and morphogenesis, among others (**Fig. 5J-K**). Codon-usage calculation revealed strong enrichment of Met-depleted genes in the up-regulated GO terms on the transcriptome and translatome levels, and vice versa (**Fig. 5L-N, Fig. S6N-P**). Given that Met is encoded by a single codon, its codon and amino acid usage bias is consistent. However, GO analysis of translation efficiency did not reveal similar terms, suggesting that the gene expression alteration for these GO terms are primarily on the transcriptome levels (**Fig. 5M-P**). Nascent translation of mitochondrial genome encoded peptides, measured after cycloheximide inhibition of cytoplasmic translation, was all reduced in both the single and double KO cells, confirming the RNA-seq and ribo-seq measurements (**Fig. 5Q**). Together, these studies revealed a correlation between biased Met codon usage and altered gene expression programs after D97/D133 KO.

To quantify the codon-biased gene expression on the transcriptome and translatome levels, we measured standard deviations of expression fold changes in KO vs. WT. Single KO affected primarily translation (**Fig. 5R**, top and middle, higher variation on the translatome level), while double KO affected both the transcriptome and translatome (**Fig. 5R**, bottom, higher variation on the transcriptome level). Double KO induced bigger changes in relative tRNA levels (**Fig. 5S**, panels 1-2). On the transcriptome level, double KO induced larger differences in codon and amino acid usage, than single KOs (panels 3-6). On the translatome level, double KO induced similar changes in codon and amino acid usage as single KOs (**Fig. 5S**, panels 7-10). Comparing variations between RNAseq and Riboseq, the single KO exerted more effects on the translational level, whereas the double KO already showed large differences on the transcriptome level, which persisted in the translatome level (**Fig. 5T**, panels 3 vs.7, 4 vs. 8, 5 vs. 9, and 6 vs. 10, and **Fig. 5T**). Together, the RNA-seq and ribo-seq in D97/D133 single and double KO HEK293 cells suggest a global reprogramming of the transcriptome and translatome (**Fig. 5U-V**). Loss of the snoRNAs resulted in defective target tRNAs, especially eMet-CAU, accompanied by reduced usage of Met and other codons in the transcriptome and translatome levels, leading to an imbalance in two competing gene expression programs: proliferation vs. differentiation/development/morphogenesis. The double KO induced larger differences on the transcriptome level, suggesting adaption of the cells to dramatically reduced levels of eMet-CAU and several other tRNAs (**Fig. 5U-V**). Comparing amino acid usage reprogramming in D97/D133 KO and previously reported stem cell differentiation model (**Fig. S6E**) ^54^ not only revealed consistent changes of multiple amino acids, but also highlighted the critical role of eMet tRNA loss in tipping the balance between the two gene expression programs after D97/D133 KO.

### Overexpression of D97/D133 target tRNAs rescues cell growth defects

To confirm that reduced tRNA activity and levels are responsible for the reduced translation of Met-enriched proteins, we infected WT and D97/D133 double KO HEK293 cells with lentiviruses expressing no insert (control) or tRNA eMet-CAT, or a mixture of 7 lentiviruses expressing eMet-CAT, Arg-CCT, Gly-TCC, Lys-CTT, Ile-TAT, Sec-TCA, and Trp-CCA (7tRNAs), which are reduced in the D97/D133 double KO (**Fig. 5C**, **Fig. 6A**). After puromycin selection, these tRNAs are expressed at higher levels (**Fig. 6B**). The overexpression of both eMet-CAT and the 7 tRNA mixture resulted in a partial rescue of the growth in KO lines (**Fig. 6C**). The increased cell proliferation WT cells after overexpression of 7 tRNAs suggest that levels of these tRNAs are limiting under normal growth conditions. To determine whether the Met-AUG codon bias is responsible for the reduced translation in D97/D133 KO cell lines, we constructed reporters (**Fig. 6D**). The insertion of 6xMet-ATG in the Fluc reporter enhanced protein synthesis relative to control Rluc in WT cells, which may have resulted from increased translation initiation (**Fig. 6E**). The insertion of 6xMet-ATG decreased protein synthesis in KO cell lines relative to WT, confirming that the D97/133 KO caused eMet-CAU tRNA defects.

**Figure 6.**
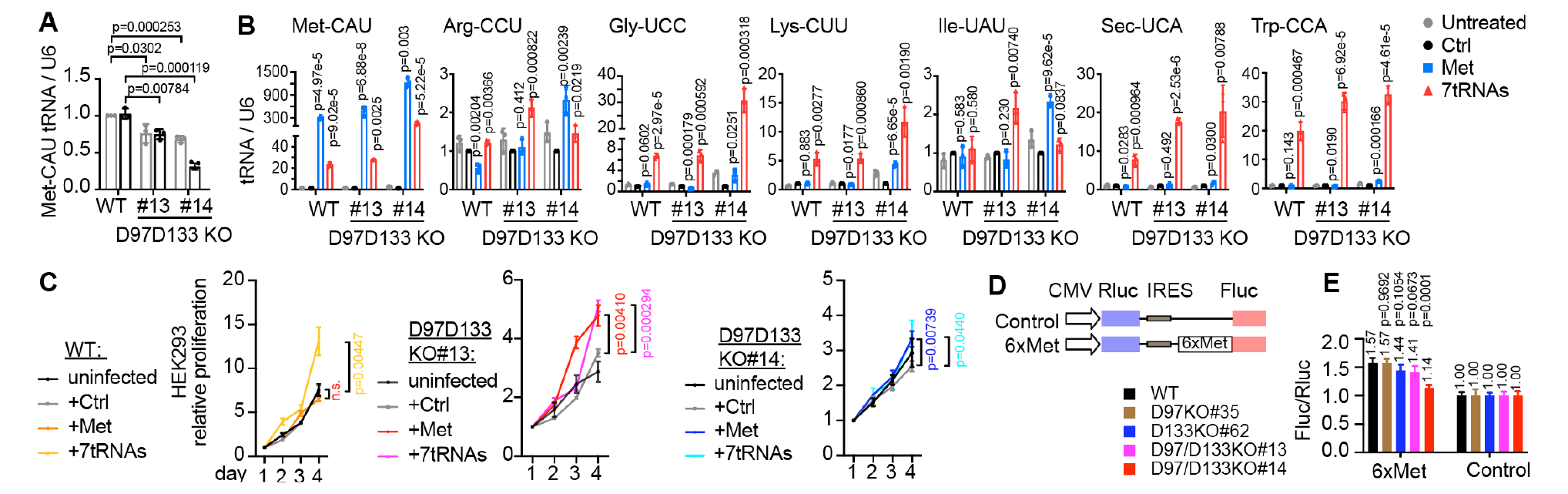
D97/D133-mediated tRNA modifications are responsible for codon-biased translation. (A) Confirmation of Met-CAU expression reduction in D97/D133 double KO cell lines, using qRT-PCR, normalized to U6 snRNA. (B) HEK293 cells were infected with pLV-EF1a-control, or pLV-EF1a expressing eMet-CAU, or mixture of pLV-EF1a expressing 7 tRNAs: eMet-CAU, Arg-CCU, Gly-UCC, Ile-UAU, Lys-CUU, Sec-UCA, Trp-CCA. After selection with puromycin for 3 days, when WT un-infected cells have died, RNA expression (qRT-PCR) and cell growth were measured for 4 days. (C) Rescue of cell growth defects by overexpressing eMet-CAU or 7 tRNAs in WT and KO cell lines. (D) Schematic of bicistronic luciferase reporter plasmids. 6xMet-ATG oligo was inserted after the start codon of Fluc. Mutagenesis oligos were inserted by PCR and cloned into pcDNA3 RLUC POLIRES FLUC vector backbone, replacing the original sequences between BamHI and BsiWI restriction sites. (E) WT and KO HEK293 cells were transfected with plasmids in panel D. After 24 hours, Fluc activity was measured to detect IRES-mediated translation, whereas Rluc activity was measured to detect cap-dependent translation, which served as an internal control. The y axis of normalized Fluc / Rluc ratio corresponding control transfected control plasmid in the same cell line. All data are representative of at least three independent experiments. p values were shown for each KO cell line relative to WT.

### Mouse snoRNAs D97 and D133 regulate stem cell states

Given the induction of development-related gene expression programs in human HEK293 cells after D97/D133 KO, we tested whether these snoRNAs regulate mES self-renewal and differentiation. The TC1 mES cells were differentiated into embryoid bodies, which contain cell types from all three germ layers (**Fig. 7A**). ASO KD of mouse Snord97 and Snord133 (D97/D133) reduced their levels by ∼50% (**Fig. 7B**). Pluripotency mRNAs for *Nanog*, *Sox2* and *Oct4* increased while markers for the three germ layers were skewed, favoring the mesoderm and endoderm (**Fig. 7C**). Contrary to D133, D97 KD does not seem to affect ectoderm fate, suggesting slightly different functions of these two snoRNAs. In particular, the cardiomyocyte (CM) *Myh6* increased after both KD (**Fig. 7D**). These results are consistent with the upregulation of genes involved in differentiation, development, and morphogenesis in HEK293 cells (**Fig. 5**). To further analyze the roles of D133 snoRNA in CM differentiation, we knocked out D133 using CRISPR (**Fig. S7A**). Cell growth slowed down significantly, similar to the D133 KO in HEK293 cells (**Fig. 7E**). We differentiated the WT and D133 KO mES cells to EBs and CMs (**Fig. 7F**). mES gross morphology and self-renewal remained the same (**Fig. S7B**), yet all KO clones significantly increased the speed and efficiency of CM formation, from ∼40% to ∼75% (**Fig. 7G-H**). The *Myh6* mRNA was induced earlier, and to higher levels throughout differentiation, in the D133 KO (**Fig. 7I**). The D133 KO also increased mRNA and protein levels of pluripotency markers, such as *Nanog, Sox2* and *Oct4*, similar to the ASO KD (**Fig. 7J-L**).

**Figure 7.**
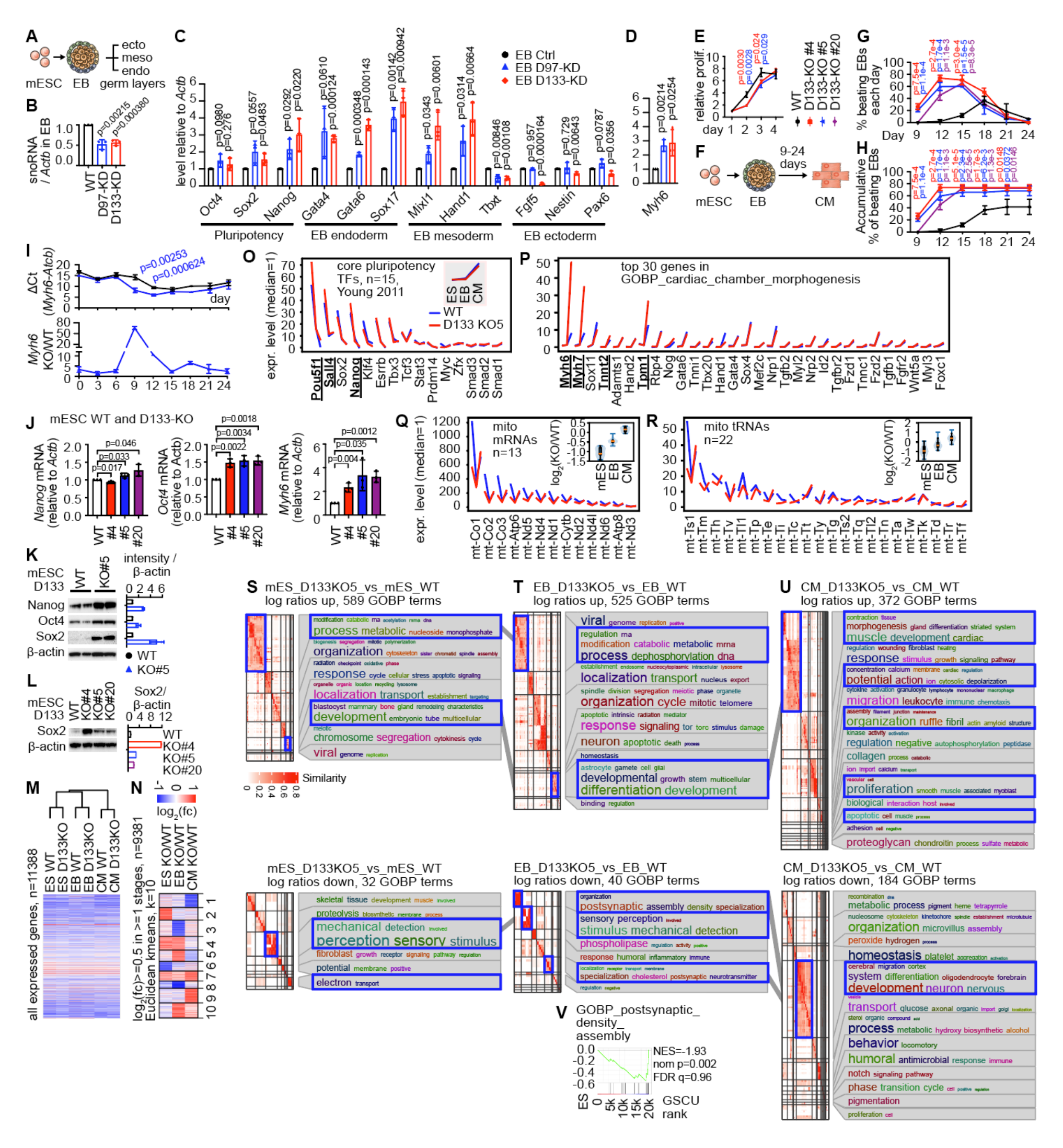
Mouse D97 and D133 regulate stem cell differentiation. (A) Diagram of mES differentiation into embryoid bodies that contain three germ layers. (B-D) After D97 and D133 ASO treatment, mES cells were differentiated to EBs. Expression of the snoRNAs (B), mRNA markers for pluripotency and germ layers (C), and cardiomyocyte mRNA Myh6 (D) from EBs were measured by qRT-PCR, and normalized to *Actb*. P values: unpaired two-sided t-tests between the KD and untreated EBs (Ctrl). (E) Proliferation of WT and D133 KO cells (clones #4 and #5). (F) Diagram for directed mES differentiation into EBs and cardiomyocytes (CM). (G) WT and D133 KO mES cells (3 clones) were differentiated into CMs. Beating CM patches were counted every 3 days. p values are from two-sided t-test, between each KO cell line and WT, and color-coded. (H) Same as panel G, except that accumulative numbers of beating CM patches are reported. (I) Expression dynamics of *Myh6* mRNA in WT and D133 KO clone #5, plotted in delta-Ct values vs. *Actb* mRNA (upper panel) or in fold difference (KO vs. WT, lower panel). (J) Expression of pluripotency and CM mRNAs in WT and D133 KO mES cells, measured by qRT-PCR. (K-L) Pluripotency factor protein levels in WT and D133 KO mES cells, measured by western blots. (M). RNA-seq data for 6 groups of samples were normalized to their medians and clustered. Biological replicates were merged before clustering. After merging replicates and removing genes with read counts <10, the samples were log-transformed, normalized for each sample and clustered using uncentered correlation complete-linkage method. (N) Log2 fold change, or log2(fc), was calculated for each gene between D133KO5 and WT. K-means clustering of the 3 stages, on genes with log2(fc)>=0.5 in at least 1 stage. The 10 groups are labeled. (O-R) Expression dynamics of core pluripotency factors (O) ^97^, top 30 genes in GOBP cardiac chamber morphogenesis (P), and all mitochondrial-encoded RNAs (Q-R), for the two genotypes across ES, EB and CM. RNA levels in were normalized so RPM median=1. Pou5f1: Oct4. In panels Q-R, stage specific expression of mitochondrial RNAs are summarized in the violin plus box plots. (S-U) Altered mRNAs based on log ratios between D133 KO clone 5 vs. WT at the ES, EB, and CM stages were median-normalized and tested for gene set enrichment using GSEA and MSigDB m5.all collection. Then enriched gene sets were clustered based on term similarity and tested for enrichment of GO terms. Example GOBP term clusters that are consistent across the 3 stages were highlighted in blue boxes and linked with gray lines. (V) Example downregulated GOBP term postsynaptic density assembly. Three biological replicates were used to report the standard deviations (error bars). P values indicate unpaired t-tests between each KO and WT mES cell line, unless otherwise noted. See also Figure S7.

To determine the molecular mechanisms driving the faster and skewed differentiation, we performed RNA-seq on WT and KO mES, EB and CM (**Fig. S7C**). We observed significant global changes in snoRNA and tRNA levels across the 3 stages (**Fig. S7D**), but the mES D133 KO did not change relative tRNA levels between KO and WT (**Fig. S7E**), consistent with the HEK293 D97/D133 single KO lines. The global expression differences among stages were bigger than between D133 KO and WT (**Fig. 7M**). In each stage, the KO induced distinct differences (**Fig. 7N**). The pluripotency TFs all dropped from ES to EB and CM stages while cardiac development related genes were induced, confirming the successful differentiation (**Fig. S7F-H**). Consistent with the qRT-PCR and western blots in D133 KD and KO, the pluripotency factors *Pou5f1* (*Oct4*), *Sall4* and *Nanog* increased after KO in the ES stage (**Fig. 7O**), whereas key factors in cardiac development increased in D133 KO vs. WT (**Fig. 7P**), also consistent with the KD results (**Fig. 7D**). Mitochondrial mRNAs and tRNAs are significantly reduced in the ES stage, but not later (**Fig. 7Q-R**). The return to normal of mitochondrial gene expression in D133 KO is consistent with the efficient differentiation to CM. Interestingly, several other mitochondrial metabolic processes, such as the one-carbon cycle, are upregulated, suggesting dysregulation of metabolites with potential roles in altering epigenetic status of D133 KO stem cells (**Fig. S7I-J**). Analysis of gene set enrichment in D133 KO vs. WT revealed consistent upregulation of development-related terms (**Fig. 7S-T**), especially cardiac development (**Fig. 7U**). In contrast, neurodevelopment related genes were down-regulated in KO vs. WT (**Fig. 7R-V**), again consistent with the D133 KD studies (**Fig. 7C**). Together, this analysis revealed enhanced pluripotency and CM gene expression programs in mouse D133 KO, consistent with the skewed and more efficient CM differentiation phenotype, and also consistent with the upregulation of GO terms in differentiation/ development/morphogenesis in HEK293 cells due to loss of D97/D133 target tRNAs (**Fig. 5U-V**).

## DISCUSSION

### Discovery and mechanistic insights into the snoRNA targetome

The large numbers of orphan snoRNAs in higher eukaryotes have remained a mystery for decades. In this study, we used several orthogonal approaches to discover and validate new snoRNA targets and modifications across multiple ncRNA types, including rRNAs, snRNAs, snoRNAs and nearly all nuclear-encoded tRNAs. For C/D snoRNA targets and Nm sites, we provided evidence from PARIS2, evolutionary conservation, optimized dRMS, CLIP (enrichment and gapped reads), and mass spectrometry. Several tRNA Nm sites seem to be sub-stoichiometric, suggesting heterogeneous tRNA populations with potentially different functions (**Fig. 2S-T** and **Fig. S3**). For H/ACA snoRNA targets, we provided evidence from PARIS2 and CLIP, but so far, we have not validated the Ψ sites in tRNAs. In addition to the recent discovery of D97/D133-guided eMet-CAU Cm34 in eukaryotes, earlier studies have predicted or validated a few sRNA-guided Nm sites in archaeal tRNAs ^3, 12–17, 93^. Remarkably, phylogenetic analysis suggested a deep evolutionary conservation of interactions that guide Met Cm34 in archaea and eukaryotes (**Fig. 2O-Q**). Even though no previous studies have found H/ACA snoRNA-guided Ψ in tRNAs, computational prediction and engineered constructs have suggested their possible existence ^94–96^.

PARIS2 not only captured snoRNA-target interactions with base pair resolution, but also provided new mechanistic insights. While earlier studies suggested these interactions typically range between 10 and 21 bps ^8^. Even though many duplexes discovered here are shorter, which may have precluded accurate computational discovery, they are still more stable than random shuffles. The frequently detected bipartite duplexes for D/D’ guides likely strengthened the stability of otherwise weak interactions (e.g., **Fig. 1S** and **Fig. 2J-N**). However, we cannot exclude the possibility that a subset of them are “biological noise” or may have functions beyond guiding modifications, as is the case for multiple well-studied snoRNAs, such as U3, U8 and U13. The detection of pre-tRNAs as snoRNA targets is further confirmed by the discovery of interactions with tRNA exon-intron junctions in archaea and plants (**Fig. 2G, Fig. S2E** and **Fig. 2O**). In eukaryotes, such interactions likely occur in the nucleolus, where snoRNAs and tRNAs are enriched ^98^. These early modification events may be prerequisites for subsequent modifications by other tRNA-targeting enzymes (**Fig. 3J-N**).

### Linking snoRNAs to tRFs, another class of regulatory RNAs

We demonstrated broad impacts of snoRNA-guided tRNA Nm sites in tRNA stability (**Fig. 4**). Earlier studies have shown that Nm stabilizes RNA duplexes ^99^. Alternatively, methylation of 2’-OH that participate in hydrogen bonding may disrupt some tertiary structures. Even though overall tRNA structures were not impacted dramatically (**Fig. 4M**), local changes are likely. Both modification chemistry and structural changes may play a role in regulating tRNA fragmentation during stress. tRFs have emerged as an important class of regulatory molecules in a variety of biological and pathological contexts ^57^. Many tRNA modifications have been shown to regulate tRNA cleavage. Our study reveals a mechanistic link between snoRNAs and tRFs, two large families of regulatory ncRNAs. Given the changes in multiple types of modifications upon snoRNP loss (either FBL KD or snoRNA KO, **Fig. 3J- N**), it remains unclear which modification defects are directly responsible for the reduced stability. Despite the functional rescue by tRNA OE (**Fig. 6**), it is likely that the increased eMet tRFs also play a role in the reprogrammed transcriptome/ translatome and skewed stem cell differentiation ^100^.

### snoRNAs controlling codon-biased gene expression programs

The D97/D133 single and double KO may reprogram the transcriptome and translatome via several distinct mechanisms. In the single KO strains, even though the tRNA levels did not change, the reduced translation of Met-enriched proteins suggests the eMet tRNAs are defective. Reduction of translation upon increased Met codon content in the reporter is consistent with this possibility (**Fig. 6D-E**). On the other hand, the double KO significantly reduced several D97/D133 target tRNAs, especially eMet-CAU, reprogramming both the transcriptome and translatome to adapt to the skewed tRNA pool. This effect is likely also dependent on active nucleases that degrade defective tRNAs. Beyond the Met-AUG codon, the transcriptome of double KO cells exhibited a remarkable dichotomy of decreasing A/U ending codons, and increasing G/C ending ones, consistent with previous reports of a dual program of translation control between proliferation vs. development, differentiation, and morphogenesis ^53, 54^. This is likely a secondary effect of the primary defects of D97/D133 target tRNAs, altered epigenome, and/or A-to-I editing. Together, our work revealed a snoRNA-controlled cellular translation economy: specific snoRNAs regulate target tRNA activity and levels – the “supply”, which influences the corresponding codon usage in mRNAs – the “demand” (**Fig. 5U-V**).

It is not a coincidence that Met-enriched gene ontologies include ribosome, spliceosome, and mitochondrial respiration, which are all essential for cell proliferation. In particular, oxidative phosphorylation related proteins encoded by both the nuclear and mitochondrial genomes have some of the highest ratios of Met residues in the proteome in higher animals ^101^. Proteins with long half-lives or close to sources of reactive oxygen species, such as those listed above, are prone to oxidative damage. Oxidized Met residues in proteins can be easily repaired by dedicated enzymes, i.e., methionine sulfoxide reductases, that exist in all three domains of life ^102^. On the other hand, the short-lived proteins involved in development, differentiation, and morphogenesis, are relatively depleted of Met. Thus, the divergent regulation of the dual program of proliferation and differentiation by the D97/D133-Met tRNA sub-network is an evolutionary inevitability. The subsequent mitochondrial metabolic reprogramming may alter key metabolites that participate in DNA, RNA and histone modifications, creating a mechanistic link between the codon-biased translation and stem cell epigenetics ^103^.

### Cell-type and developmental-stage specific functions of snoRNPs

A recent study observed no growth or overall translation defects in either single or double KO of the HAP1 cell line, a near-haploid derivative of KBM-7 that came from a chronic myeloid leukemia (CML) patient ^15^. In contrast, in both human HEK293 (embryonic adrenal medulla, a neuro-ectodermal lineage) ^104^ and mouse ES KO cells, we observed significantly slower growth and translation, as well as codon-biased gene expression on the transcriptome and/or translatome levels (**Figs. 5** and **7**). These differences likely reflected cell-type and developmental stage specificity of snoRNA and tRNA expression (e.g., **Fig. S7D**), as well as their functions^4, 105^. Consistent with this, we observed differential effects of D133 KO on gene expression during differentiation of mES to EB and CM (**Fig. 7N-U, Fig. S7I-J**). The skewed differentiation potential of mES to the three germ layers further suggested the divergent effects of these snoRNAs in different lineages (**Fig. 7C**). Further studies are needed to understand the precise defects of tRNAs and cell-type specificity of snoRNA functions.

### Limitations of the study

Several technical challenges have impeded the discovery of snoRNA targets and their modifications. Many snoRNAs are expressed at very low levels or in a tissue specific manner ^4^. Current UV and psoralen crosslinking methods remain inefficient and biased ^65^, especially for H/ACA snoRNAs (**Fig. S4**). The short snoRNA guides and our incomplete knowledge of the interaction mechanisms preclude accurate computational prediction. Even though global tRNA Nm levels reduced significantly after FBL KD, we were only able to validate a small number of them using dRMS (**Tables S2, S3** and **S6**), despite improved performance (**Fig. S1A-J**). In addition, high throughput analysis of pseudouridine is challenging, making it difficult to validate the H/ACA snoRNA target sites in tRNAs ^106^. The hierarchical snoRNA-tRNA-mRNA networks and their roles in codon-biased cellular state control remains poorly defined. The pleiotropy of snoRNAs in targeting multiple tRNAs and other ncRNAs, the redundancy of multiple snoRNAs with identical or similar target sites, and the multi-copy tRNA gene families are challenging for genetic analysis. The interdependency of tRNA modifications results in complex defects (**Fig. 3J-N**), masking the direct effects of each genetic perturbation. Even though we observed earlier and more beating CMs, their functionality and potential application in regenerative medicine require further characterization.

## STAR METHODS

Detailed methods are provided in the online version of this paper and include the following:

## RESOURCE AVAILABILITY

Lead contact

Materials availability

Data and code availability

## EXPERIMENTAL MODEL AND SUBJECT DETAILS

Cell lines

## METHOD DETAILS

Design of biotinylated antisense oligos

snoRNAs enrichment and PARIS2 library construction

Chromosome-associated RNA (caRNA) extraction

Optimized dRMS for small RNAs

RNA knockdown studies

CRISPR/Cas9-mediated snoRNA knockout

LC-MS (ESI) detection of RNA modifications

Oxidative stress treatment

tRNA in vitro angiogenin (ANG) cleavage assay

Melting curve analysis

Lentiviral overexpression of snoRNAs and tRNAs

Cell proliferation assay

Bulk RNA-seq library preparation

Ribosome profiling (ribo-seq) library preparation

HPG labeling for monitoring global protein synthesis

Detection of mitochondrial-encoded nascent peptides

Luciferase Reporter Assay

Mouse embryonic stem cell (mES) differentiation

Western blot

## QUANTIFICATION AND STATISTICAL ANALYSIS

snoRNA conservation analysis

snoRNA target predication

PARIS2 data analysis

RiboMethSeq data analysis

RNA structure and conservation analysis

Human PAR-CLIP and eCLIP data analysis

Yeast CLIP data analysis

Bulk RNA-seq data analysis

Ribo-seq data analysis

Gene set enrichment analysis

## SUPPLEMENTAL INFORMATION

Supplemental information can be found online.

## ACKNOWLEDGEMENTS

We thank members of the Lu lab for discussion. We thank K. Machida for antibodies, O. Bell for the TC1 cell line, and the Albany RNA Epitranscriptomics and Proteomics Resource for mass spectrometry. The Lu lab is supported by startup funds from the University of Southern California, NHGRI (R00HG009662 and R01HG012928), NIGMS (R35GM143068), USC Research Center for Liver Disease (P30DK48522), Illumina and USC Keck Genomics Platform (KGP) Core Lab Partnership Program, USC ALPD (P50AA011999), the Norris Comprehensive Cancer Center (P30CA014089) and USC Center for Advanced Research Computing.

## AUTHOR CONTRIBUTIONS

Z.L. conceived and designed the project. M.Z. performed PARIS2 analysis of snoRNA targets, CLIP analysis, human cell line KO and KD studies. K.L. and J.B. performed reporter assays, rescue experiments, and functional studies of mES KO cell lines. M.Z. and R.v.D. developed the dRMS method. W.Z. and J.F.C. prepared iPS-derived cell lines. M.Z., R.v.D., M.A. and B.S.L. performed MS analysis of modifications. M.Z. and Z.L. wrote the software for data analysis. M. Z. K.L. J.B. and Z.L. performed sequencing data analysis. Z.L. wrote the manuscript with input from all other authors. Z.L. supervised the project.

---

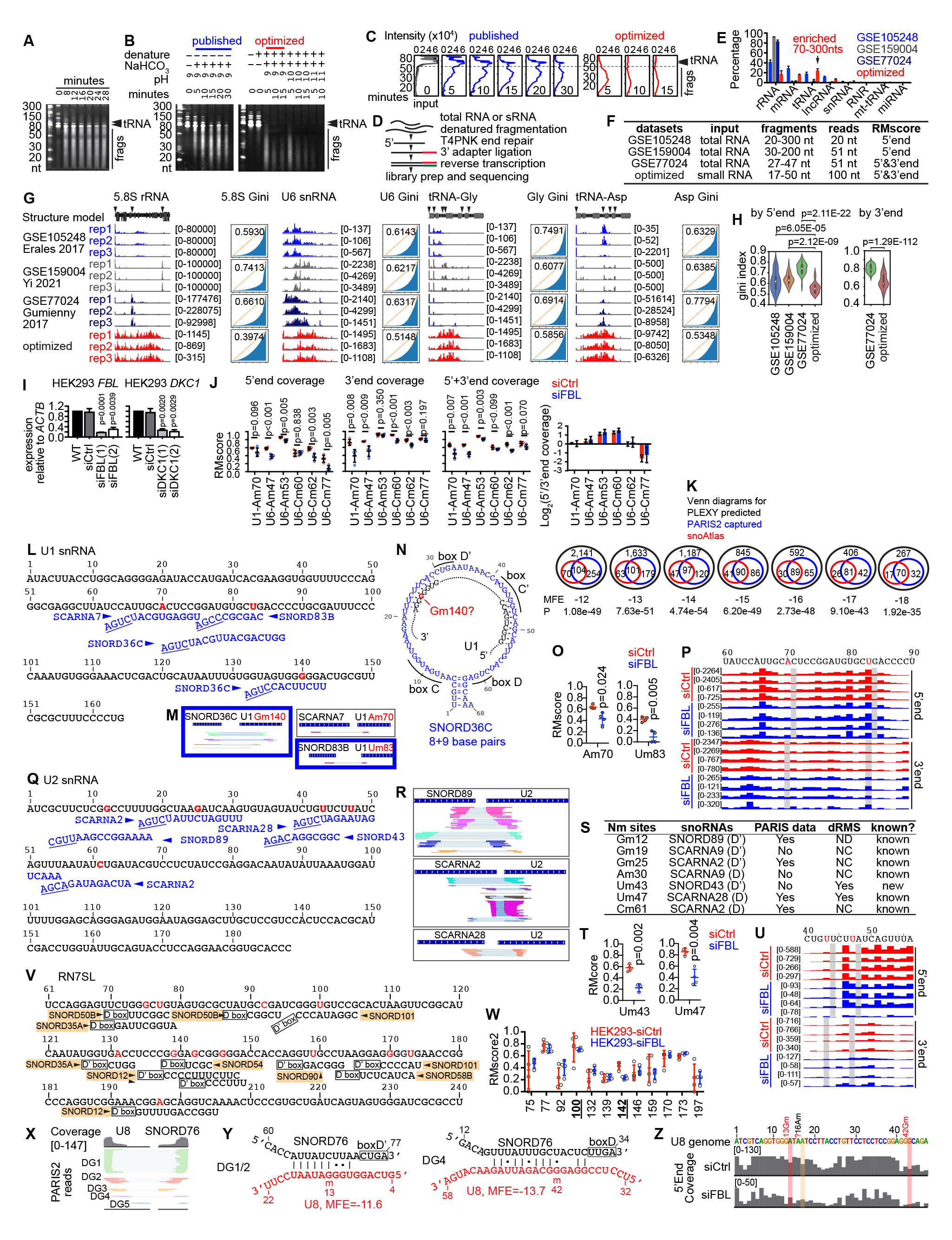

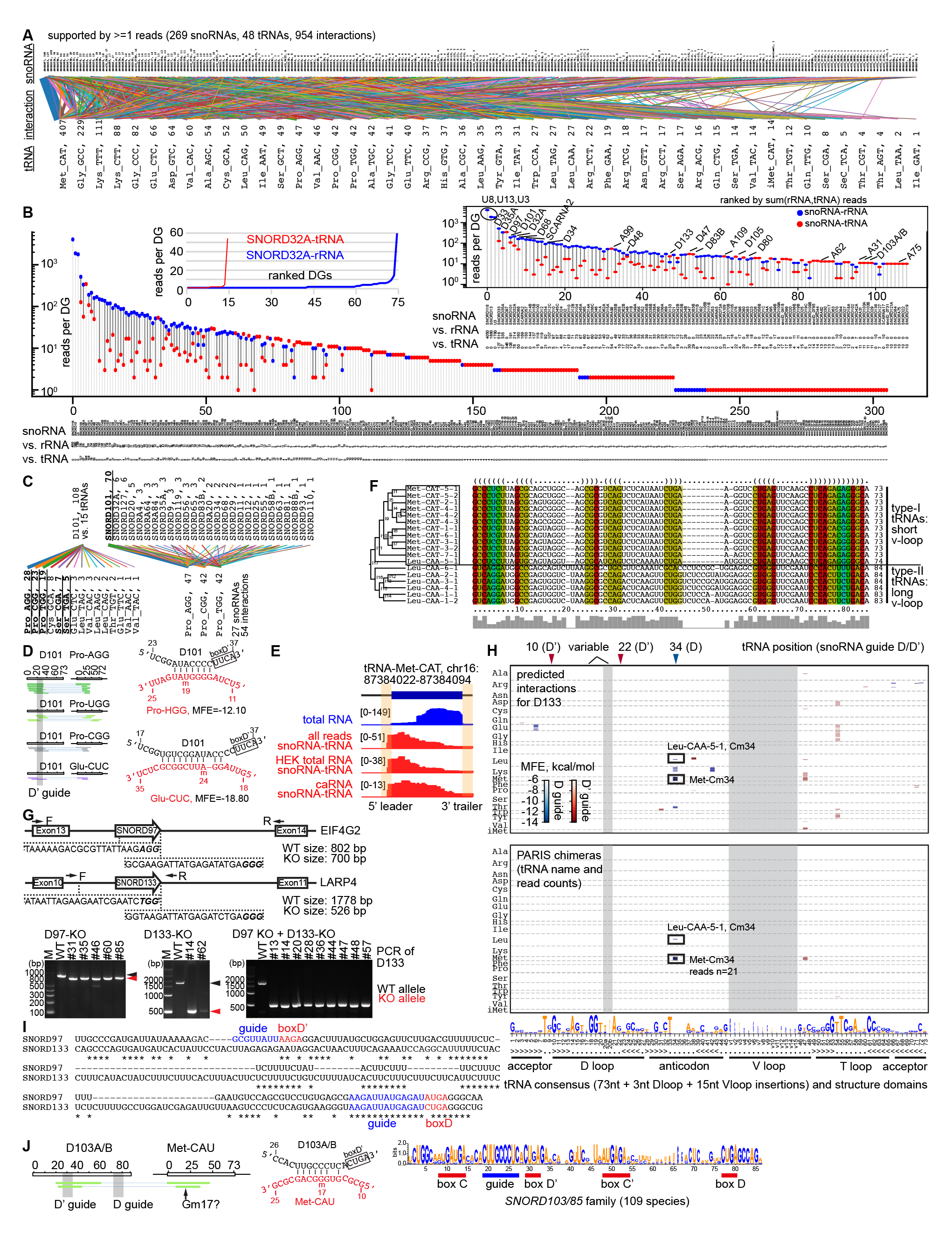
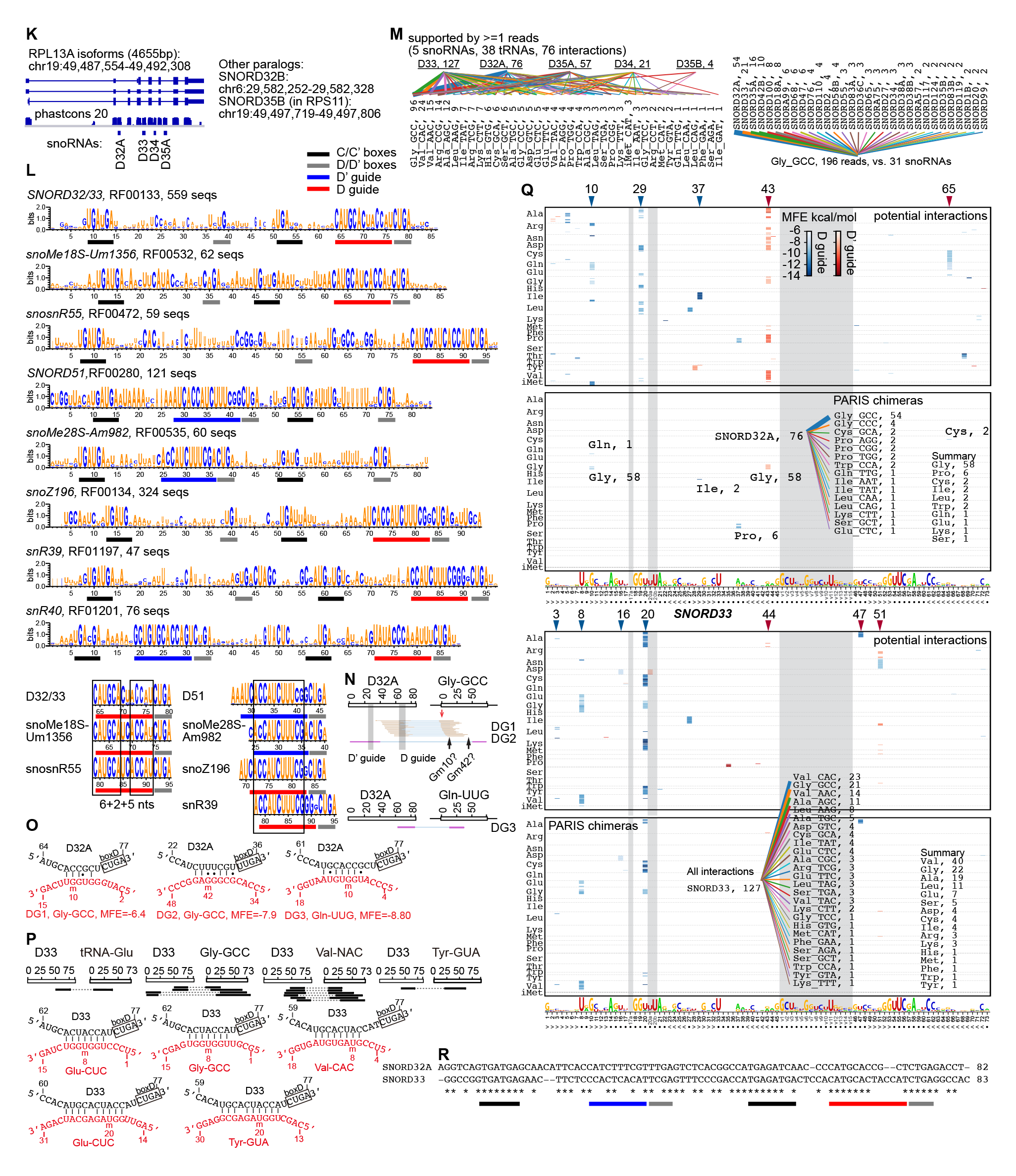
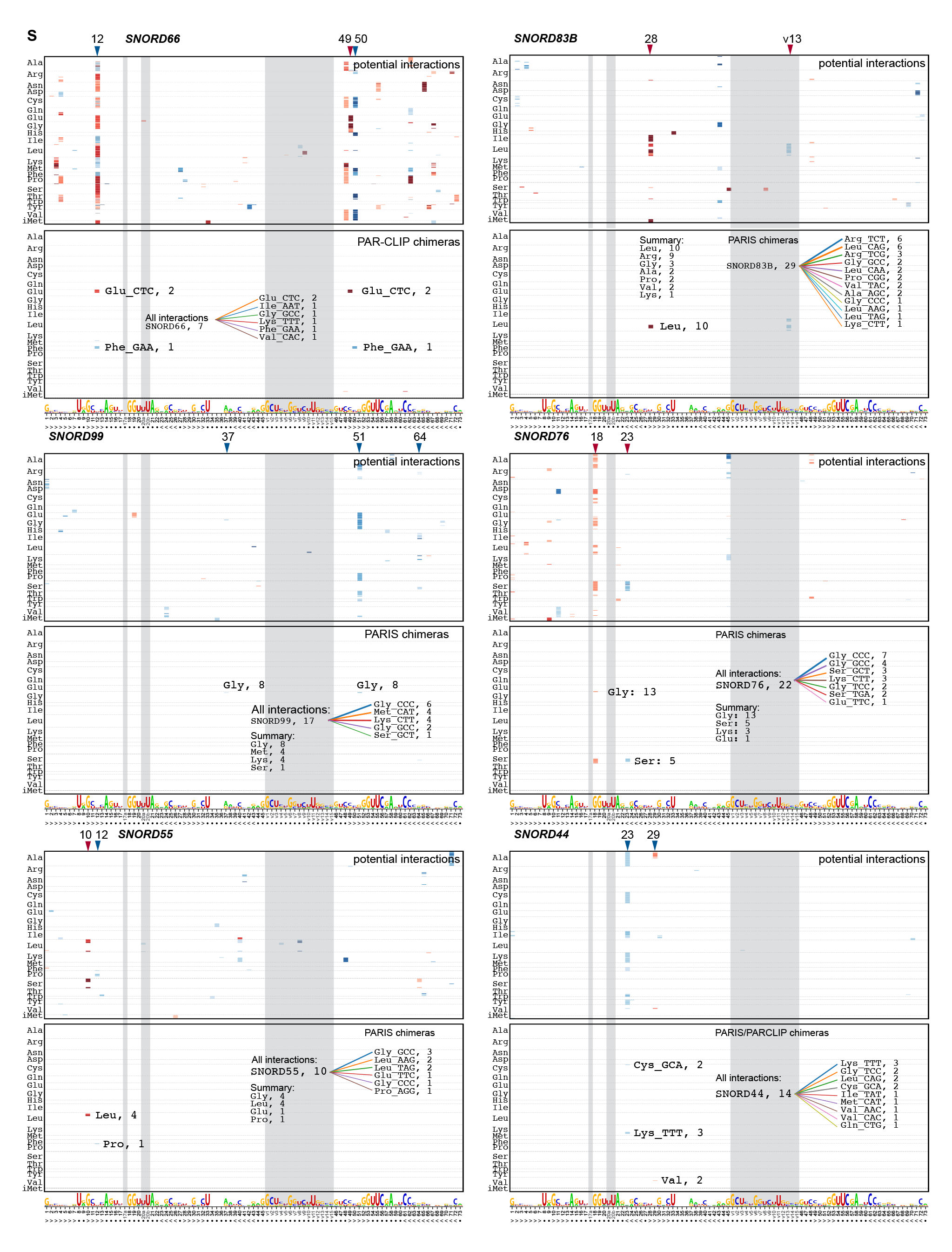
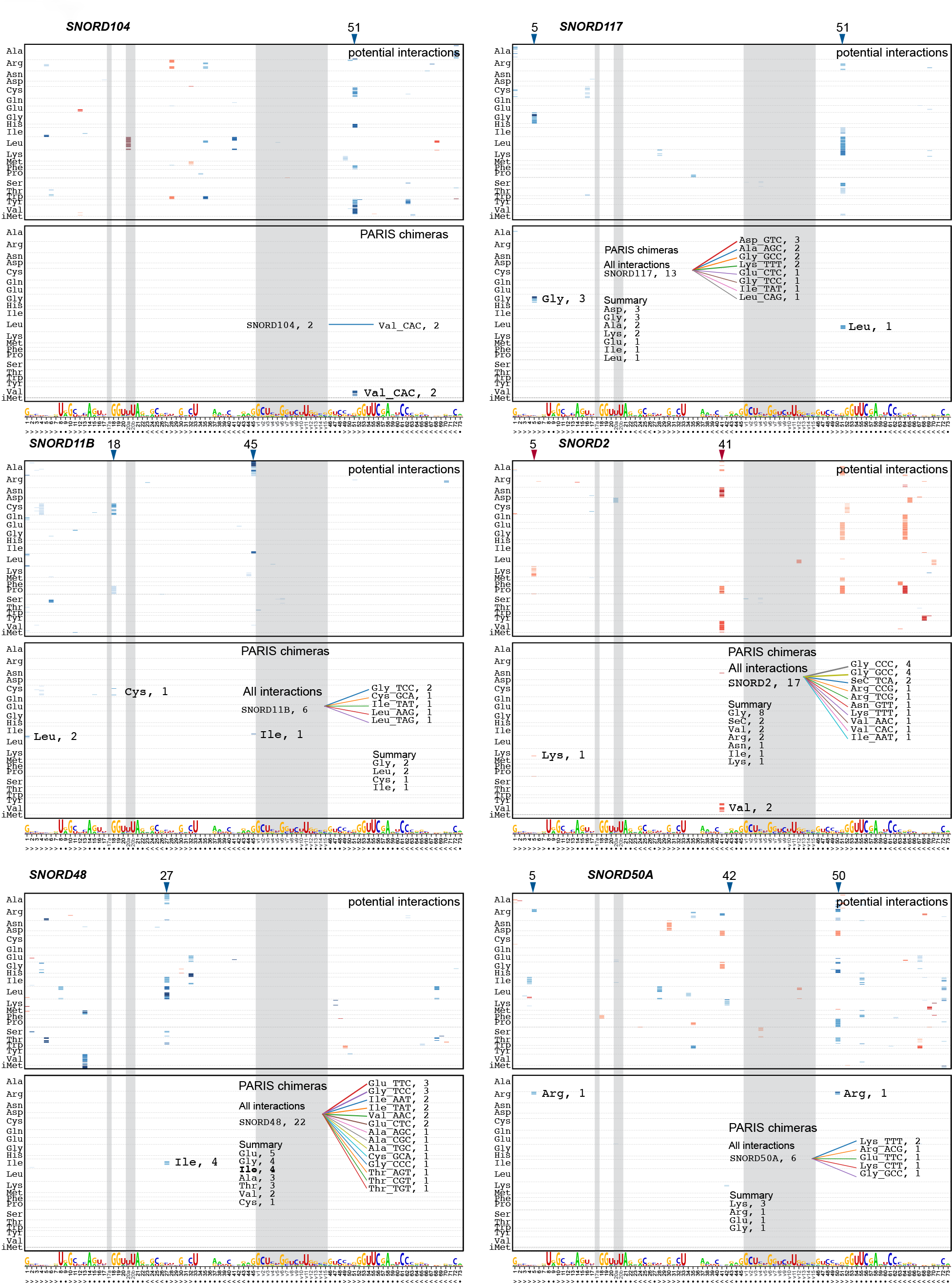
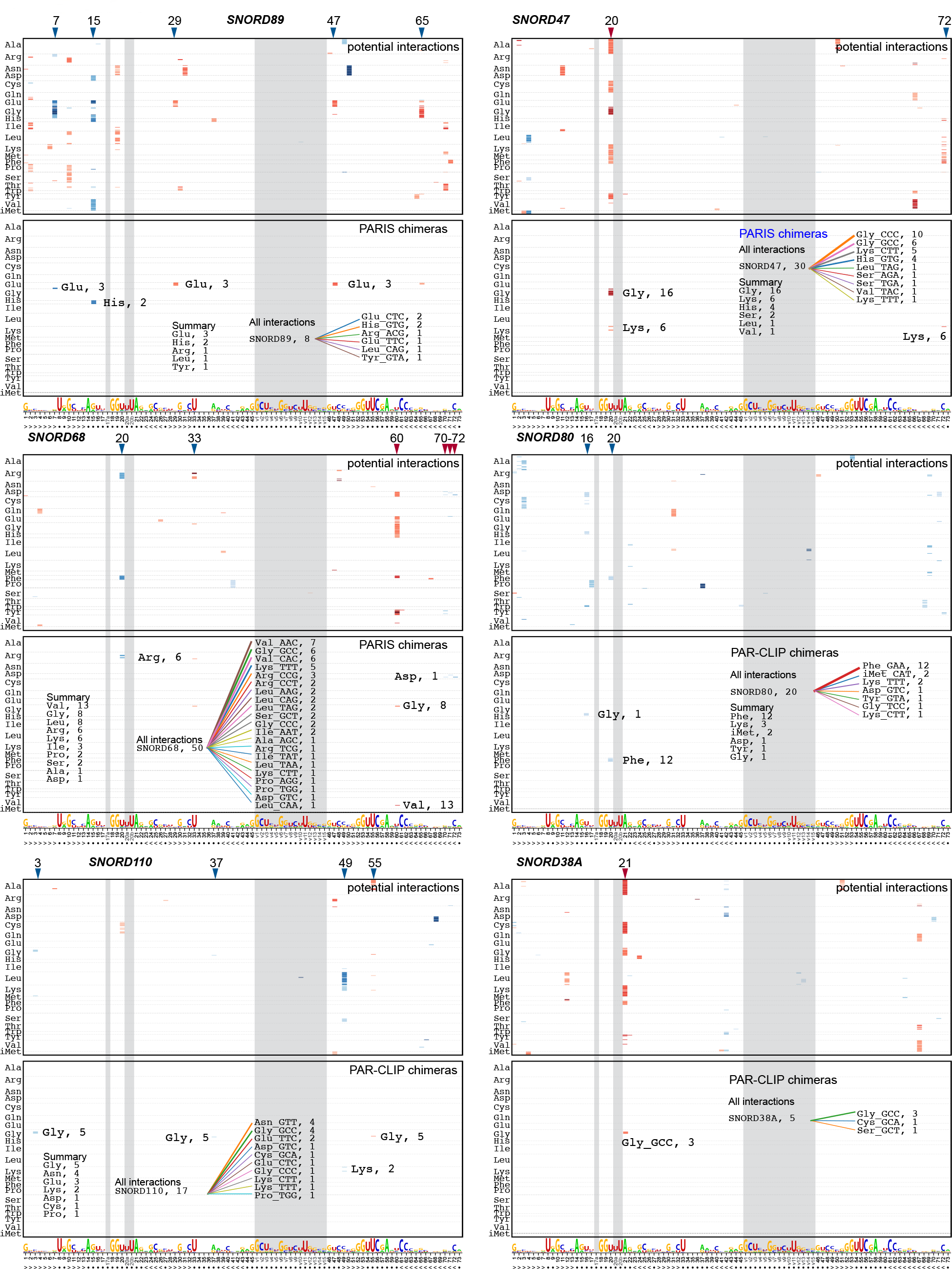
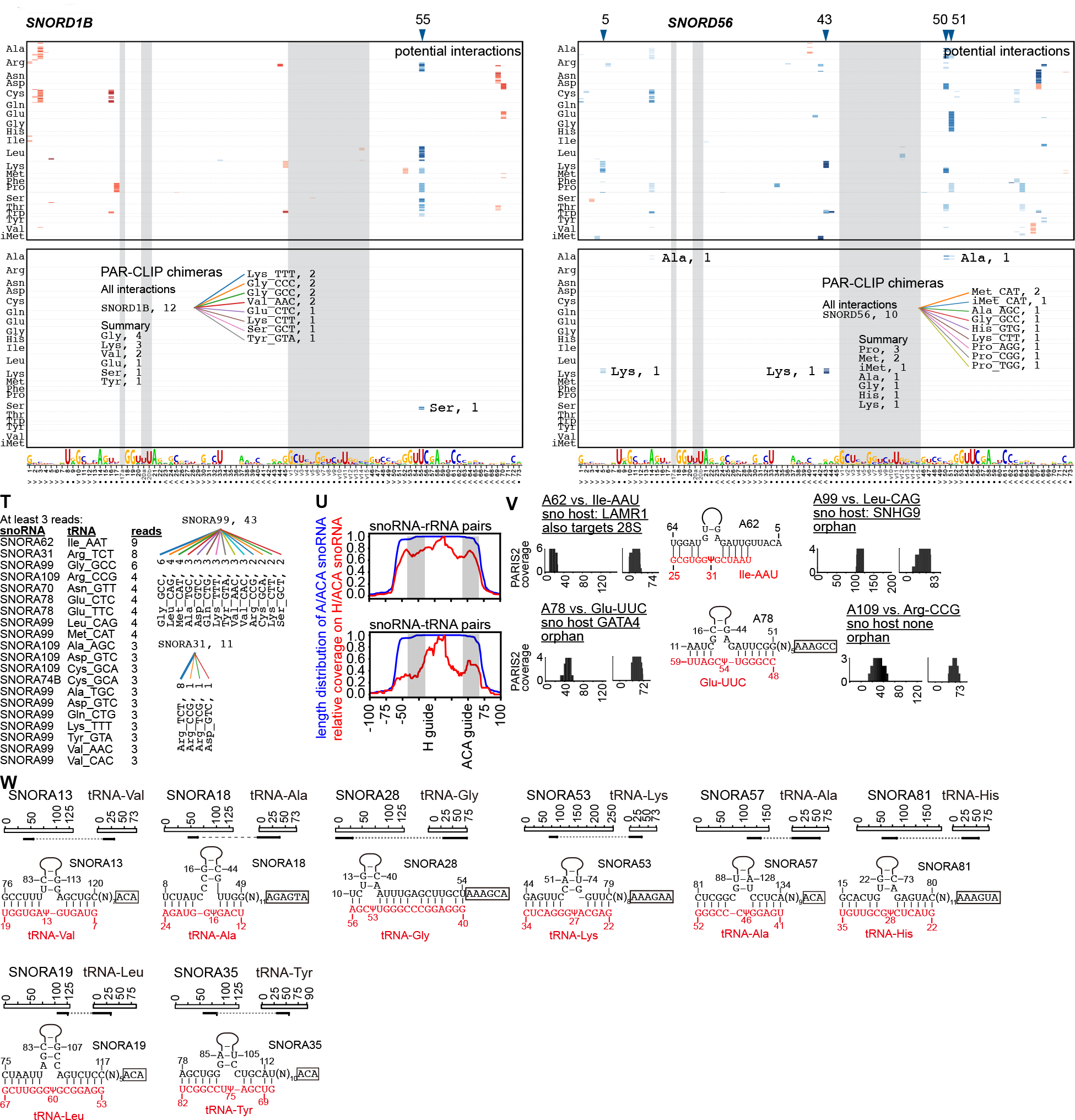

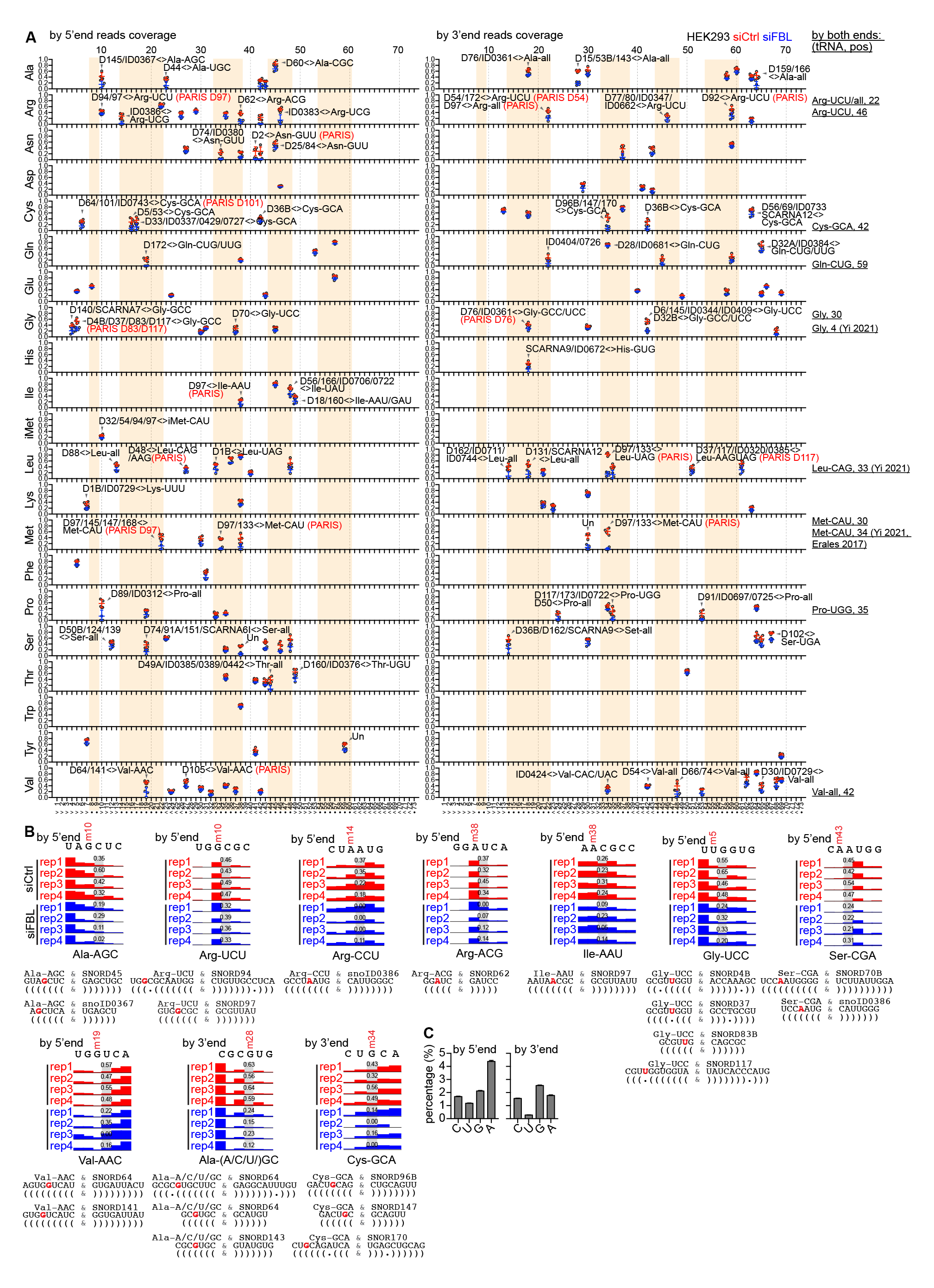
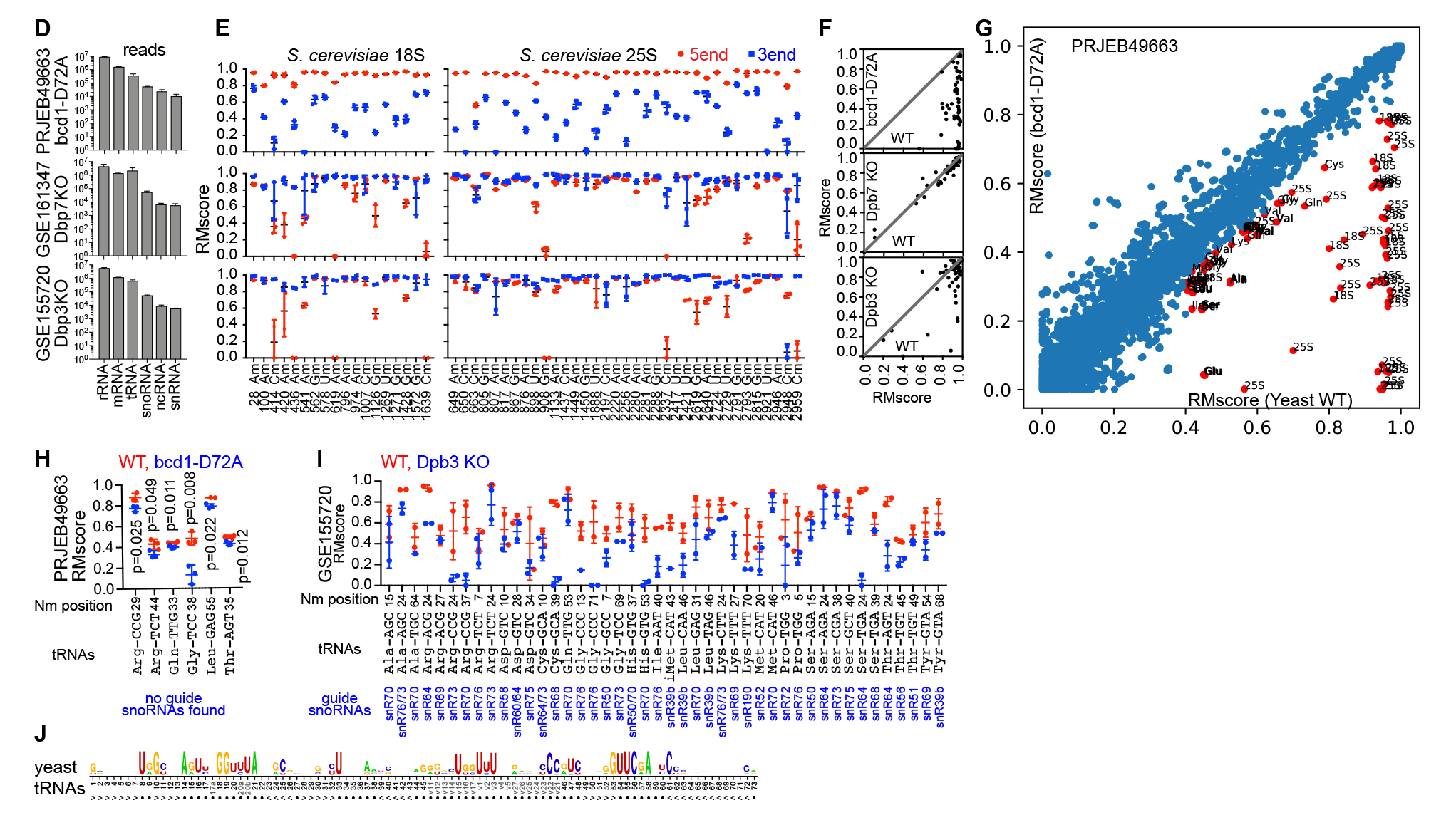

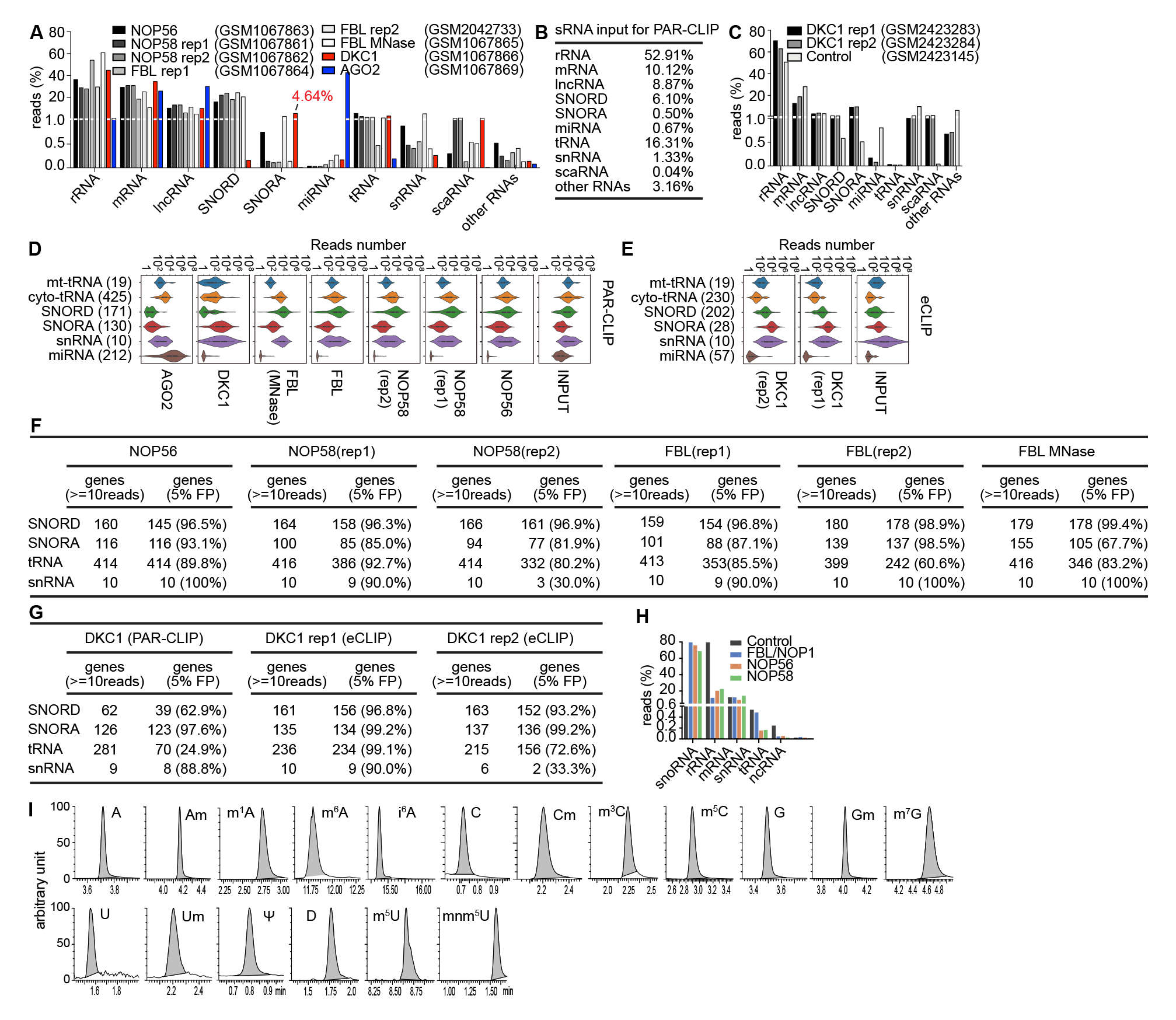

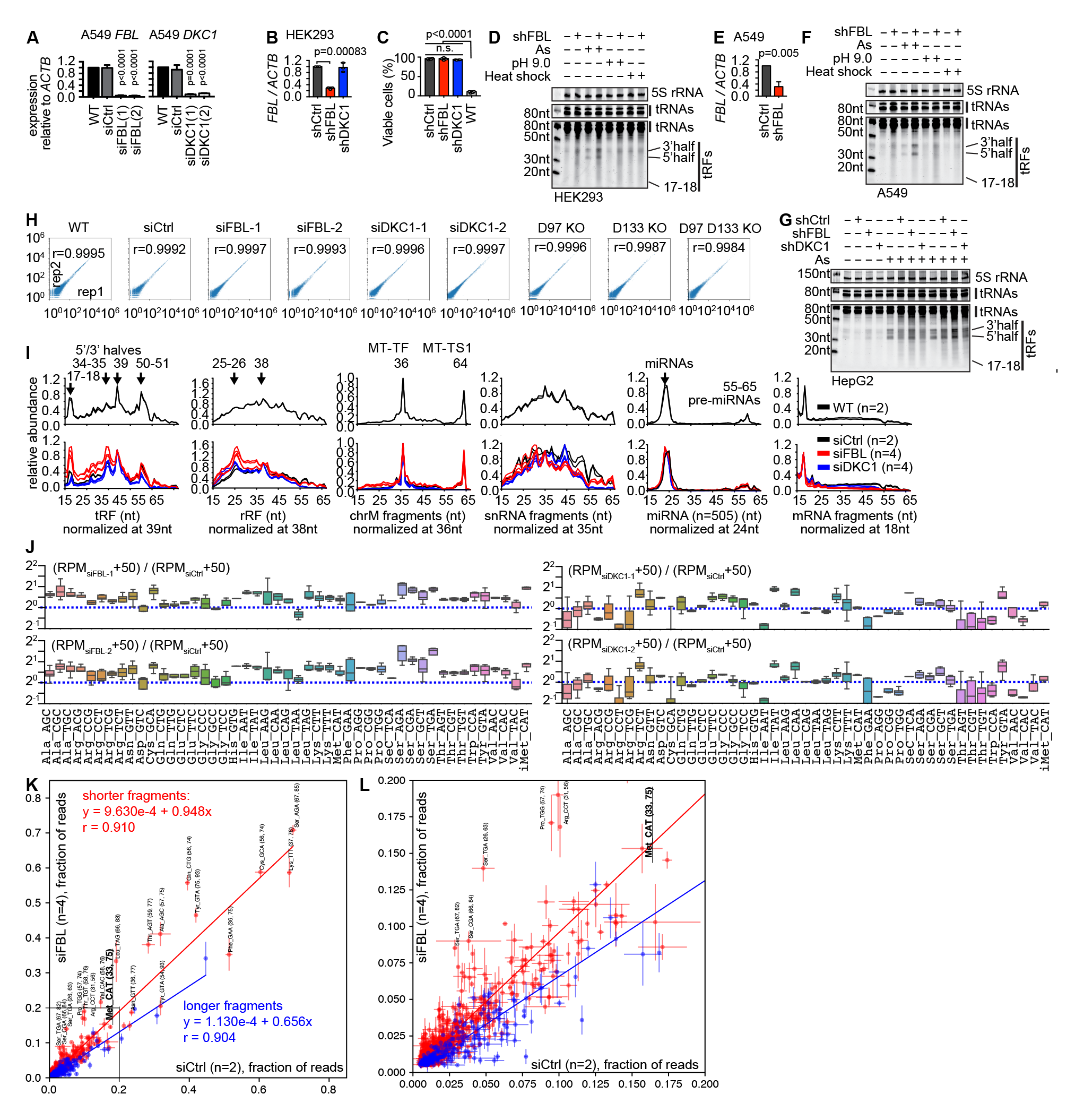

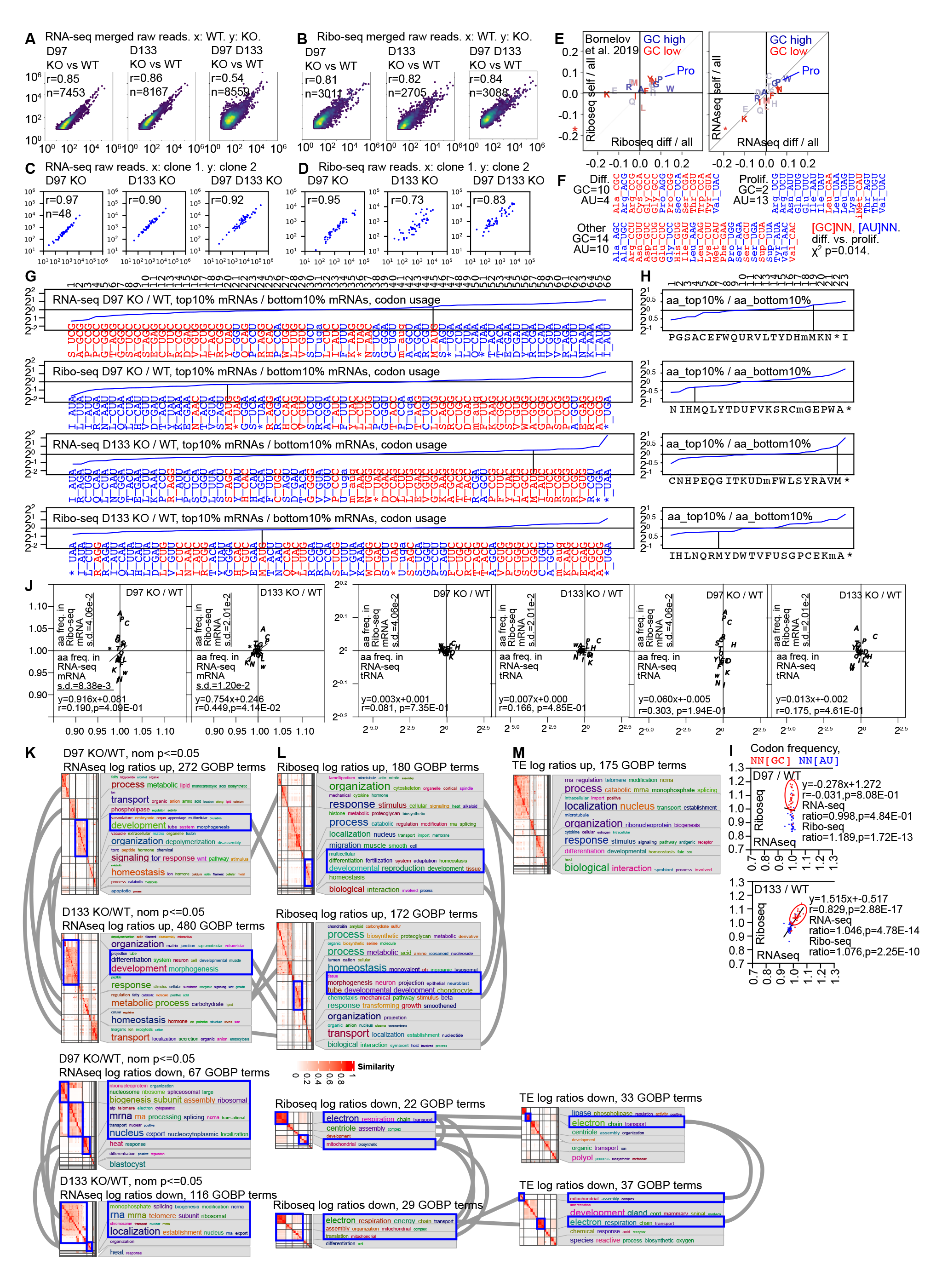
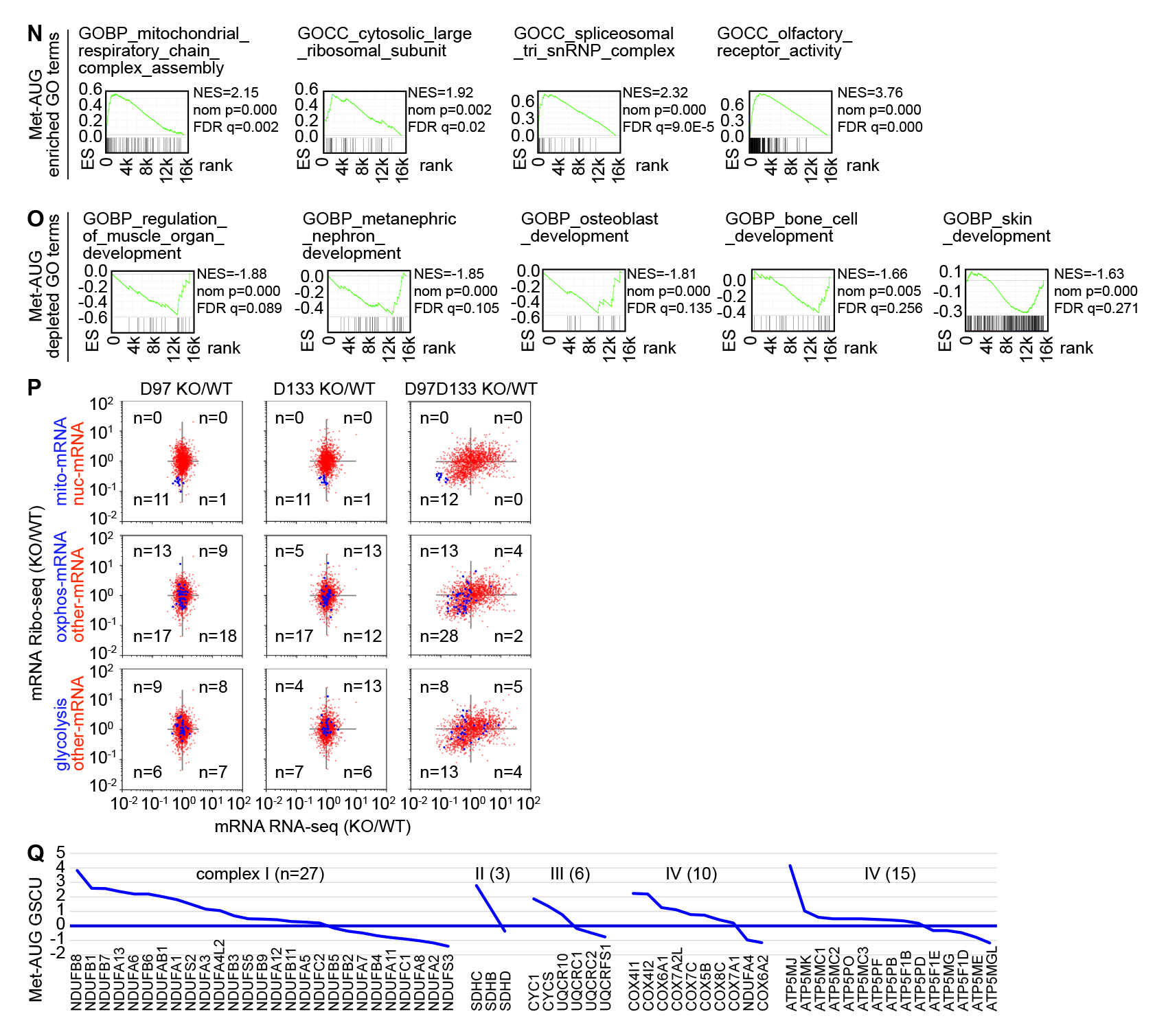

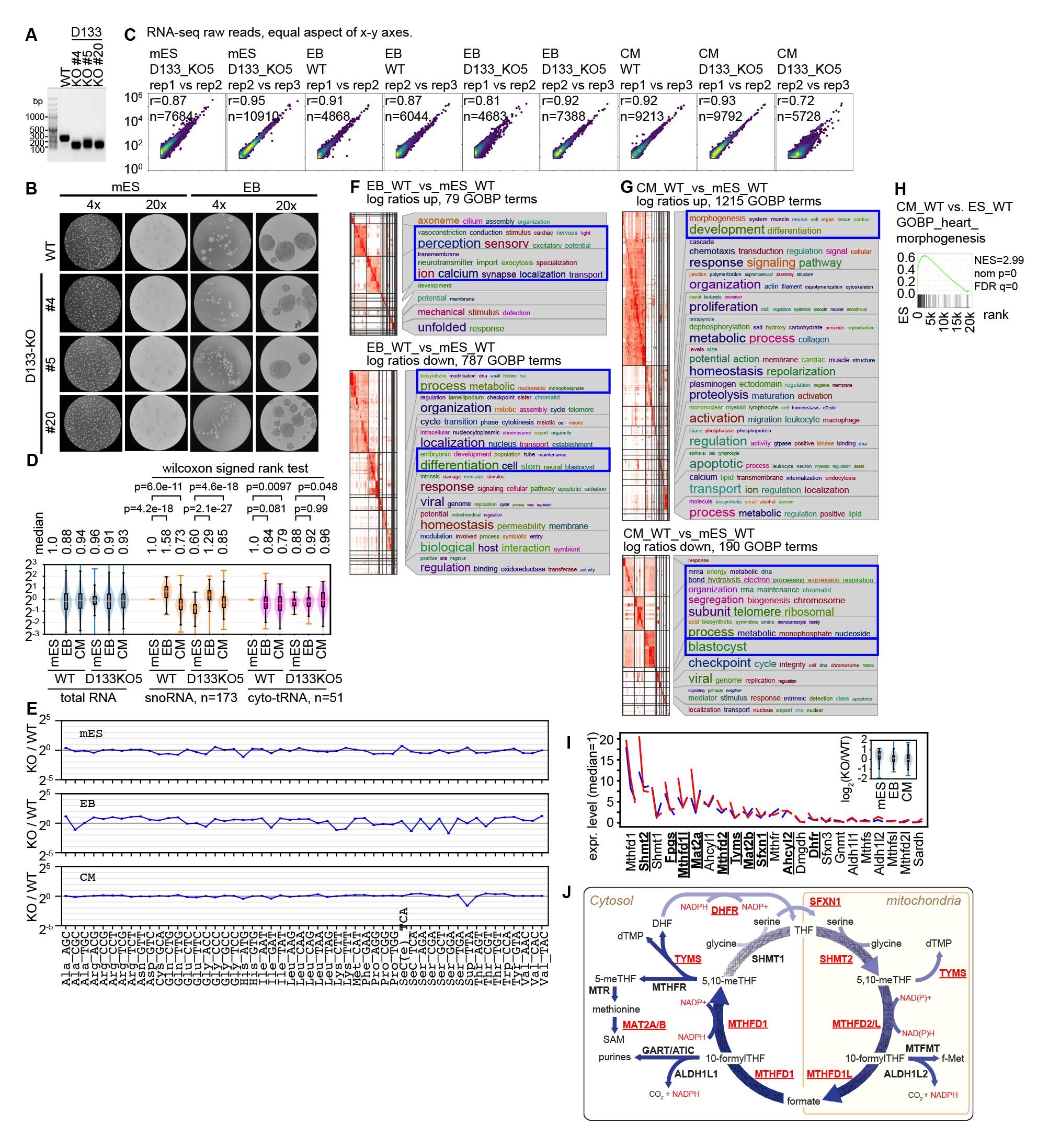

