## Supplemental figure legends for "snoRNA-guided tRNA 2’-O-methylation controls codon-biased gene expression and cellular states"

**Figure S1. Development of dRMS and discovery of new snoRNA targets.** dRMS was optimized to reduce salt concentration and include ~50% DMSO for denaturation and tested on snRNAs after FBL KD (panels A-K). Integrating PARIS2 and dRMS, here we confirm known and discover new Nm sites in human U1 (L-P), U2 (Q-U), RN7SL (V-W) and U8 (X-Z).
(A) Adding formamide during alkaline hydrolysis does not enhance fragmentation. Hydrolysis conditions used here: to 10µL purified small RNAs (sRNA, 70-300nt) at 10ng/µL in 95% formamide, add 10µL 100mM bicarbonate buffer at pH 9. Incubate at 95C for 0-28 mins, precipitate RNA and separate 50ng RNA in a 12% urea-PGAE gel. The 70-80nt tRNA bands remain after 28 mins hydrolysis. Other abundant ncRNAs in the range of 100-300nts also persisted after more than 10mins, e.g., 5S/5.8S rRNAs, spliceosomal snRNAs, snoRNAs and RN7SL.
(B) Small RNA (sRNA) fragmentation optimization. Hydrolysis of 100 ng sRNA (70-300 nt) was performed in 10 mM sodium bicarbonate buffer at different pH for different times. Denaturation: 100 ng of sRNA was denatured in 95% DMSO solutions for 3 mins at 98C prior to addition of the bicarbonate buffers. Compared to the published RMS [1], inclusion of DMSO in hydrolysis resulted in complete loss of full length tRNAs and other abundant ncRNAs above tRNAs.
(C) Quantification of sRNA fragments from the published and optimized conditions at pH 9.
(D) dRMS pipeline. On top of the standard RMS method, reduced salt and ~50% DMSO is used during fragmentation.
(E) Summary of RMS datasets compared in this study, including Erales et al. 2017 (HeLa cells, GSE105248 and PRJEB43738) [2], Gumienny et al. 2017 (HEK293 cells, GSE77024) [3], and Yi et al. 2021 (C4-2 PCa cells, GSE159004) [4]. Percentage of reads mapped to each category of transcripts are listed. The optimized dRMS method was applied to enriched sRNAs in the range of 70-300, while previous published datasets were from total RNAs.
(F) Summary of RMS data properties compared in this study. When read lengths are shorter than fragment lengths, or when RT enzymes that have terminal transferase activities are used, only the 5' ends are useful for calculation of RNA 2'-O-methylation scores (RMscore for short).
(G) 5'end reads coverage on 5.8S rRNA, U6 and tRNAs, as well as calculated Gini indices for these transcripts. Compared to published fragmentation conditions, our optimized dRMS method generates more uniform start positions of fragments (lower Gini index values). Incomplete fragmentation leads to biased ends near the transcript starts, i.e., more full-length transcripts, e.g., panels A-B, or near internal loops of the RNA structures. In the dot-bracket structure models, dots indicate single stranded regions, whereas brackets indicate base pairs.
(H) Violin plot showing the Gini index for 5' and 3'end read coverage of published RMS and our optimized dRMS methods. In the box plots, whiskers represent the max and min. The top and bottom of the box represent the 1st and 3rd quartiles. The open circle is the median. tRNA loci with sequencing read coverage more than 3000 in all three datasets were analyzed in here. For the 5'end and 3'end reads coverage, 55 and 341 tRNA loci were analyzed, respectively.
(I) Relative mRNA levels of FBL and DKC1 in HEK293 cells after FBL/DKC1 KD using two different sets of siRNAs. Data are mean ± s.d. of n=3 independent experiments. P values indicate two-tailed, unpaired t-test between each KD and the siCtrl (siRNAs that do not target human mRNAs). WT indicates untreated cells.
(J) Analysis of known Nm sites in snRNAs from dRMS data in HEK293 cells, comparing siRNA KD of FBL vs. control. n=4 for each condition. p values are from two-sided unpaired t-tests. RMscore was calculated using 5'end and 3'end read coverage separately or together. The end coverage of known Nm sites with ± 2nt reads were shown. Rules to call Nm sites: p value <= 0.05, mean coverage of (Nm sites ± 2nt) >= 20, and, for specific Nm sites, if p value <= 0.05, we use the end with higher sequencing coverage to call Nm levels. For example, if 5'end coverage >= 3'end coverage, we will use 5'end reads coverage to calculate Nm level. This analysis also suggests unavoidable systemic errors in RMS data, even though both can reveal partial Nm loss for most sites after FBL KD. For each Nm site, the coverage of ends for the 5' and 3' are compared and calculated in log2 scale. Some sites have more 5' ends, while other sites have more 3' ends. This variation in coverage between the two ends affects accuracy of RMscore calculation.
(K) Venn diagrams for overlaps of PARIS2 captured and snoAtlas annotated snoRNA-rRNA interactions within the PLEXY predicted set at various MFE cutoffs (kcal/mol). P values are from hypergeometric tests.
(L) U1 has 3 reported Nm sites, Am1, Um2 and Am70, the last one of which is guided by SCARNA7 (U90) [5]. PARIS2 and dRMS revealed several potential new sites. Guide snoRNAs are colored blue. D/D' box elements are underlined.
(M) PARIS2 gapped reads supporting three snoRNA-U1 interactions. In particular, new interactions are highlighted in blue boxes. Both SNORD36C and SNORD83B were previously predicted to guide modifications on rRNAs (http://snoatlas.bioinf.uni-leipzig.de/) [6]. Their interactions with U1 suggest functional pleiotropy.
(N) Secondary structure model for the bipartite SNORD36C-U1 interaction (8 and 9 base pairs). The putative Gm140 site is indicated. The interaction with D guide is strong (9 base pairs) but too far from the D box.
(O) Methylation levels measured by dRMS on two sites, Am70 and Um83. Gm140 was not detected by dRMS despite the strong interaction data from PARIS2. Four biological replicates were included for the dRMS analysis.
(P) The dRMS reads end coverage profiles on Am70 and Um83 were plotted for both 5' and 3' ends.
(Q) U2 has at least 10 validated Nm sites. A subset of the known and potential new Nm sites on U2 snRNA are shown here [5].
(R) Example known snoRNA-U2 interactions captured by PARIS2.
(S) List of Nm sites with PARIS2 and dRMS support. ND: not detected. NC: no change. Am1, Um2, Gm11 and Cm40 are not included. Only Nm sites with either PARIS2 or dRMS support were shown in the diagram.
(T-U) Methylation levels for two Nm sites based on dRMS. The 5' and 3' reads end coverage profiles were plotted separately (U).
(V) PARIS2 and dRMS identified potential Nm sites on RN7SL. Predicted target sites between the RN7SL RNA and snoRNAs with at least 1 PARIS read support for the interactions are shown here. D/D' box motifs are omitted.
(W) Only 5' end dRMS data were used to calculate RMscore. Only Nm levels of Um100 and Gm142 were significantly reduced after FBL KD (two-sided t-test p<0.05).
(X) Strong interaction between snoRNAs U8 (SNORD118), which is mutated in the neurodegenerative disease Leukoencephalopathy with Calcifications and Cysts (LCC) [7], and SNORD76, which was previously shown to target the 28S rRNA.
(Y) Secondary structure models of the two SNORD76-U8 interactions, and the MFE values (kcal/mol). Two sites are potentially modified. Gm13 is in the 5' end of U8, which forms multiple interactions with itself, as well as with the 28S rRNA [8,9]. The ~30nts at the 5' end of U8 is required for 28S rRNA processing [10], and frequently mutated in LCC [7].
(Z) dRMS data for the U8 snoRNA, including sites Gm13 and Gm42. Replicates were merged due to low coverage. The potential guide snoRNAs for Am16 is unknown at this moment.

**Figure S2. Characterization of the snoRNA-tRNA network**. This figure includes details for the following snoRNAs: the global network (A-B), D101 that binds both rRNAs and tRNAs (C-D), D97/D133 and eMet-CAU related interactions (E-J), RPL13A snoRNAs and related interactions (K-R), other C/D snoRNAs (S), and a subset of H/ACA snoRNAs (T-W). In addition to PARIS2 data, we also provided additional evidence derived from evolutionary conservation and structure modeling.
(A) The global network of snoRNA-tRNA interactions, including all connections supported by >=1 reads. snoRNAs and tRNAs are ranked by total numbers of gapped reads (PARIS + CLIP) connecting them to their partners. The line thickness is scaled to the square root of read number.
(B) Numbers of reads supporting snoRNA-rRNA (blue dots) and snoRNA-tRNA (red dots) interactions compared side by side for each snoRNA. Interactions were ranked by the sum of numbers for both types of interactions. Zoom-in view shows interactions supported by at least 10 total reads. The 3 snoRNAs that guide rRNA processing, U3 (SNORD3 paralogs), U8 (SNORD118), and U13 (SNORD13) have the strongest interactions with rRNAs. Some of the snoRNAs only bind rRNAs, e.g., U8, U13, U3, SCARNA2; some of them only bind tRNAs, e.g., D97, A99, D47, D83B; the rest bind both, e.g., the D33, D32A, D34, D35A cluster. Example D32A interactions with both rRNAs and tRNAs were ranked by reads per DG (interaction). The strongest interactions with rRNAs and tRNAs have similar numbers of read support.
(C) All 15 tRNA targets of SNORD101 and the numbers of reads supporting these interactions (left). All 27 snoRNA partners of tRNA-Pro and the numbers of reads supporting these interactions (right). All but one nucleotide among the 3 Pro tRNAs is different, which is in the anticodon wobble position.
(D) Alignments of PARIS2 reads supporting D101 interactions with Pro and Glu tRNAs, highlighting the consistent binding to the conserved D’ guide, but not the variable D guide region (left, see alignments in Fig. 1). Structure model of the D101 interactions with Pro-HGG and Glu-CUC (right). The D’ box element is inconsistent with the CUGA consensus. In HGG, H stands for A, C or U.
(E) All reads mapped to eMet-CAU tRNA (blue) and snoRNA-tRNA interaction reads (red). Reads supporting snoRNA-tRNA interactions have 5’ and 3’ end extensions, suggesting that they occur on the tRNA precursors.
(F) Human Leu-CAA-5-1, the only type-I gene among Leu-CAA copies, clusters with eMet-CAU genes, suggesting that it originated from mutation of an ancestral eMet-CAU gene, likely U37-->A (CAU to CAA). All Leu-CAA-1/2/3/4 genes have introns, which are removed before alignment. All human Leu-CAA and eMet-CAU genes were aligned using the default setting of LocARNA. Leu-CAA and Leu-UAA tRNAs are modified at position 34 as well (Leu-UAA: ncm5Um34, Leu-CAA: hm5Cm34, Fig. 1A in previous study [11]).
(G) Construction of D97/D133 KO HEK293 cells. EIF4G2 hosts only D97. LARP4 hosts only D133. Open arrows represent snoRNAs. Open rectangles represent neighbor exons in host genes. The forward (F) and reverse (R) primers are used for PCR validation. The genomic sequences targeted by sgRNAs are shown in sense orientation. The PAM (or complementary) sequences are in bold italics. PCR of genomic DNA from snoRNA KO HEK293 cells were visualized on 2% agarose gels. The double KO was made from D97 KO.
(H) Heatmap for PLEXY predicted (upper panel) and PARIS2/CLIP discovered (lower) Nm sites guided by D133. Each row represents a tRNA gene, grouped by anti-codon. Shades of blue represent D guide targets, whereas shades of red represent D’ guide targets. All tRNAs are aligned to a standard model of 73nts plus insertions at the D-loop (not to be confused with the snoRNA D box/guide) and T-loop. Three most targeted sites by D97 are labeled here as a reference: 10, 22 and 34.
(I) Alignment of human D97 and D133. The C, C’, D and D’ boxes (red), putative antisense elements (blue) are highlighted. Asterisks mark identical nucleotides.
(J) PARIS2 reads supporting D103A/B interaction with eMet-CAU mapped to the D’ but not D guide, potentially guiding Gm17 modification. Structure model of the D103A/B interaction with eMet-CAU, mediated by the D’ guide is consistent with evolutionary conservation of the D’ guide. D103/D85 family homologs were identified from Rfam, aligned using MAFFT and visualized in WebLogo. Positions with deletions above 70% were removed from the consensus. The D’ but not D guide is conserved in evolution.
(K) Location and conservation (UCSC phastcons 20) of RPL13A snoRNAs. Two paralogs are not in RPL13A: D32B and D35B.
(L) Alignment of the D32/33 family sequences and related snoRNAs. Homologs were identified from Rfam and aligned using MAFFT. The alignment was visualized using WebLogo. D32 and D33 belong to the same family RF00133 in Rfam, which is part of a large clan CL00054 that contains 8 families: snoMe18S-Um1356 (Rfam RF00532), snoMe28S-Am982 (Rfam RF00535), SNORD32/33 (Rfam RF00133), SNORD51 (Rfam RF00280), snosnR55 (Rfam RF00472), snoZ196 (Rfam RF00134), snR39 (Rfam RF01197), snR40 (Rfam RF01201). snoMe18S-Um1356 (RF00532) was annotated primarily in insects. Only one of the homologs was identified from a fungus, Smittium culicis, which is a parasite of insects. SNORD32/33 (RF00133) was annotated in both metazoa and plants. snosnR55 was annotated primarily in yeast. Together, these data suggest that this family is probably conserved in all eukaryotes. The mRNA interaction that Elliott et al. reported was with the D32A D' guide [12], and is not conserved in evolution. Analysis of guide sequences in the CL00054 clan reveals 2 groups, where SNORD32/33, snoMe18S-Um1356 and snosnR55 have nearly identical D' guide sequences, while SNORD51, snoMe28S-Am982, snoZ196 and snR39 have nearly identical guide sequences.
(M) RPL13A snoRNAs interact with tRNAs (left side), in addition to their reported rRNA targets. The gapped reads supporting the interactions are ranked. In this group, only the ones in RPL13A are expressed and interacting with targets at high enough levels to be detected by PARIS2. The Gly-GCC snoRNA-tRNA interaction subnetwork is highlighted (right side, 31 snoRNAs), where three of the RPL13A snoRNAs are ranked in the top (D32A, D33 and D35A).
(N-O) Example PARIS2 reads supporting interactions between D32 and two tRNAs (N), as well as secondary structure models (O), and minimal free energy levels (MFE, kcal/mol). It is likely both interactions with Gly-GCC occur at the same time, even though only the D guide is conserved. The D guide that binds tRNAs is the same as for the rRNA interaction.
(P) Example PARIS2 reads supporting interactions between D33 and several tRNA targets (upper panels), as well as secondary structure models (lower panels).
(Q) PLEXY predicted (potential interactions) and PARIS2-derived (PARIS2 chimeras) D32A and D33 target tRNAs. Major target sites are labeled with arrowheads. Each row presents a tRNA locus. Consensus of human nuclear-encoded tRNA alignments is shown at the bottom.
(R) Alignments of human D32A and D33 sequences. Even though these two snoRNAs have very similar D guide sequences, they are not the same, as shown by the 2nt shift.
(S) tRNA targets for other snoRNAs D33, D83B, D99, D76, D55, D44, D104, D117, D11B, D2, D48, D50, D89, D47, D48, D80, D1B, D56, D66. For each example, the upper half shows predicted pairwise interactions based on minimal free energy (MFE). The lower

half shows the pairwise interactions captured by PARIS2 and/or PAR-CLIP, and the interaction network, e.g., numbers of reads connecting the snoRNA and the tRNA. The arrows point to major target sites. The human cytoplasmic tRNA consensus sequence and structure is shown at the bottom as a weblogo.
(T) PARIS2 also identified H/ACA snoRNA-tRNA interactions, but not as efficiently as for C/D box snoRNAs. This could be due to lower expression levels, transiency of interactions, inaccessible crosslinking sites and other unknown factors. Example sub-networks are shown for SNORA99 (supported by at least 2 reads), and SNORD31, recently identified to harbor genetic variants that impair cortical neuron-intrinsic immunity to the virus HSV-1 and underlie herpes simplex encephalitis [13].
(U) For all gapped reads with one arm mapped to the H/ACA snoRNAs, the coverage was averaged over the snoRNAs. For the meta-gene analysis, the position before the H motif is set to 0. Data are normalized to max=1. For all H/ACA snoRNA-target pairs, coverage of the arms mapped to snoRNAs are summed up in red. snoRNAs targeting rRNAs are shown on the top, while snoRNAs targeting tRNAs are shown on the bottom. Blue lines are the average length distribution of snoRNAs (x axis: nucleotides). Positions for the average guide sequences are labeled.
(V-W) Examples of H/ACA snoRNA-tRNA interactions supported by PARIS2 reads. SNORA62, also known as E2, is located in an intron of RPSA, together with SNORA6. SNORA62 was previously shown to guide 18S rRNA processing and pseudouridylation of residues U3830 and U3832 in 28S rRNA [14-18]. For two snoRNAs A78 and A28, their targets are the at the TΨC loop. Pus10 and/or DKC1/Cbf5 in various organisms have been shown to catalyze this Ψ [19], but no guide snoRNAs have been reported. Earlier studies suggested that archaeal Cbf5, the DKC1 homolog can modify tRNAs, particularly pos 55, in the absence of H/ACA guide snoRNAs [19]. It has been suggested that DKC1 evolved the ability to use H/ACA snoRNAs later. In panel V, the tight clustering of gapped reads provides strong support for the validity of these interactions.

**Figure S3. Analysis of snoRNA-guided Nm sites on human and yeast tRNAs.**
(A) RNA samples (n=4) from siCtrl and siFBL HEK293 samples were collected 7 days after KD. sRNAs (50-500nt) were cut and purified from 8% PAGE gel, and Nm levels were detected by dRMS. The following abbreviations were used in the figure: D, SNORD. ID, snoID (new snoRNAs from snoAtlas database). Un, unkown. Orange highlighted regions are the loops. Significantly reduced modification levels based on 5' and 3' read ends were plotted separately, and ones that are supported by both are labeled on the right side, including further support by published data (Erales 2017 and Yi 2021) [2,4]. RMscores from the 5' and 3' ends are not always consistent due to sequencing biases, as described in Fig. S1.
(B) A subset of the snoRNA-tRNA interactions and dRMS values are described. Note: while the 3' ends report the modification status of the specified nucleotide, 5' ends report the modification status of the upstream nucleotide.
(C) Percentage of nucleotides with Nm in tRNAs, as measured by dRMS. Note this value is not directly comparable to LC/MS data of total tRNA Nm levels.
(D) Mapping statistics of three yeast RMS datasets. Bcd1 encodes an essential factor in snoRNP assembly. Loss of Bcd1 resulted in greater reduction of Nm levels in rRNAs than KD of Fbl/Nop1. The bcd1-D72A mutation causes cells to have low steady-state levels of box C/D snoRNAs, resulting in significant loss of Nm levels [20]. The Dbp3 RNA helicase participate in snoRNA processing and recycling [21]. The Dbp7 RNA helicase is required for snoRNA-dependent ribosome assembly [22].
(E) RMscore of 53 known Nm sites on yeast 18S and 25S rRNAs. For PRJEB49663, only 5'end coverage can be used. For GSE161347 and GSE155720, both 5'end and 3'end can be used.
(F) Comparison of RMscores of 53 known Nm sites on yeast 18S and 25S rRNAs between WT and mutants. Bcd1-D72A mutant has the biggest impact, whereas Dbp7 deletion has the smallest.
(G-H) Comparison of RMscores for all nucleotides in yeast rRNAs and tRNAs between WT and bcd1-D72A (G). In budding yeast, snRNAs are pseudouridylated, but 2′-O-methylation has not been observed [23], and also not observed here. Six bcd1-dependent example Nm sites in tRNAs are shown (H). p values are from two-sided t-tests.
(I) Example Nm sites on yeast 18S and 25S rRNAs between WT and mutants ($p < 0.05$, two-sided t-tests). Predicted guide snoRNA are shown at the bottom. Most yeast snoRNAs have been reported to guide rRNA modifications, and not considered orphans.
(J) Aligned human and yeast tRNA sequences, the variable loop is different between human and yeast tRNAs.

**Figure S4. Characterization of snoRNP CLIP datasets and RNA modification LC/MS data.**
(A) Bar plot of the percentages of human RNA types in PAR-CLIP data (GSE43666). The percentages of C/D box snoRNAs (SNORD) in NOP56, NOP58 (rep1), NOP58 (rep2), FBL, FBL (MNase) were 14.80%, 20.31%, 21.95%, 17.40%, 20.26% respectively. However, percentages of H/ACA box snoRNA (SNORA) in DKC1 sample was only 4.64%, suggesting the DKC1 PAR-CLIP was not as efficient as the SNORD CLIP experiments. The percentages of reads containing T to C mutation in NOP56, NOP58 (rep1), NOP58 (rep2), FBL, FBL (MNase), DKC1 were 32.56%, 64.46%, 67.03%, 47.11%, 32.81%, 6.62% respectively. The much lower T to C mutation rate for DKC1 further confirms the low efficiency of DKC1 PAR-CLIP, compared to the C/D snoRNP crosslinking results. 18S and 28S rRNAs, 7SK, and 7SL are also targets of snoRNPs but too big to be included in the 20-200nt sRNA-seq (GSM1067868), therefore their enrichment ratios are not accurate. Note, this input control dataset is not the 18-30nt small degraded (sd)RNA (GSM1067867).
(B) Percentages of reads mapped to various RNA categories in the sRNA PAR-CLIP input dataset.
(C) Bar plots of the percentages of human RNA types in DKC1 eCLIP and size-matched control datasets.
(D-E) Reads number of different types of RNAs in snoRNP PAR-CLIP (GSE43666, HEK293 cells) and eCLIP (GSE91645, HepG2 cells) datasets. For each type of RNA, only RNA loci with RPM > 1 in input samples were analyzed. The number of genomic loci of each RNA type was showed in following brackets. The data were visualized in Log10 scale in violin plots.
(F-G) For each CLIP experiment, the numbers of genes enriched above the FP cutoff were listed, and percentages were calculated relative to numbers of genes detected with >=10 reads.
(H) Percentages of reads mapped to each category of yeast RNAs.
(I) Example liquid chromatography (LC) peak profiles for nucleotides detected in the LC/MS analysis of tRNAs and rRNAs.

**Figure S5. FBL and snoRNAs protect tRNAs from degradation.**
(A) The relative mRNA levels of FBL and DKC1 in A549 cells after FBL/DKC1 KD using two different siRNAs, analyzed by qRT-PCR, normalized against *ACTB* mRNA. Data are mean ± s.d. of n=3 independent experiments. P values indicate two-tailed, unpaired t-test between each KD and the siCtrl. WT is untreated cells. siCtrl is an siRNA that does not target any human RNA.

(B) Levels of FBL and DKC1 mRNAs after stable short hairpin (sh)RNA KD in HEK293 cells, analyzed by qRT-PCR, normalized against *ACTB* mRNA. Analysis performed 5 days after 1 µg/ml doxycycline induction. Data are mean ± s.d. of n=3 independent experiments. P values are from two-tailed, unpaired t-test.
(C) shRNA KD of FBL and DKC1 did not reduce cell viability. Two days after induction of FBL and DKC1 shRNAs by 1 µg/ml doxycycline (stably integrated into genome), cells were treated with 2 µg/ml puromycin, which only kills cells without the puromycin-resistance shRNA vector. Wide type and shCtrl HEK293 cells served as control. Cell viability was measured using trypan blue staining after 5 days doxycyline induction.
(D) RNA fragment levels after various types of stress and FBL KD by shRNA. Arsenite stress: 250 µM arsenite for 4 hours; pH 9.0: 100mM Tris (pH9.0) for 4 hours; Heat shock: 65C for 15 mins.
(E) Same as panel A, except this analysis was performed using shRNAs on A549 cells.
(F) Same as panel D, except this analysis was performed using shRNAs on A549 cells.
(G) RNA fragment levels after arsenite (As) treatment and FBL/DKC1 shRNA KD in HepG2 cells.
(H) For each experimental condition, including untreated cells (wildtype, WT), mock siRNA KD (siCtrl), FBL and DKC1 KD (two different siRNAs for each mRNA), D97/D133 single and double KO, reads mapped to each gene from the two replicates of small RNA-seq data were plotted. Pearson's correlation coefficients were calculated.
(I) Fragment length distribution from different classes of RNAs, including tRNAs (tRF), rRNAs (rRF), mitochondrial derived RNAs (chrM), snRNAs, miRNAs and mRNAs, normalized to 1 at the indicated lengths (e.g., 39 for tRFs). For tRFs, major peaks are 17-18nt fragments from cleavage at the D or T loops, 34-35 and 39 nt peaks cleaved at the anticodon loop (5' and 3' halves, respectively), 50-51nt peaks representing RT stops at $m^{2,2}$G26 (50+26=76, based on the most common tRNA length: 73 nt plus 3' end CCA). siFBL promotes shorter fragments: 17-18nts, and 34-35nts. rRFs are also shorter after FBL KD. The 36nt peak MT-TF is likely a fragment from mitochondrial phenylalanine tRNA. The MT-TS1 peak is the full length 64nt mitochondrial serine tRNA1. The miRNA analysis was performed on the confident set of 505 miRNA precursors, based on the miRBase collection. The major peaks are 22nt mature miRNAs and 55-65nt pre-miRNAs. The 18nt peak in mRNAs are mis-annotated mRNAs that overlap tRNA genes, e.g., chr17:8,226,619-8,226,696, from Ser-AGA (hg38 genome). Additional 18nt mRNA artifacts include the following positions: chr17:8226607 (n=6019 reads), chr11:60756987 (n=3723), chr15:96282921 (n=2426), chr1:179095412 (n=2301), chr7:102973619 (n=2270), chr16:30756395 (n=2011), chr11:96341447 (n=1093), etc.
(J) Changes of 15-50nt RNA levels upon FBL and DKC1 KD by two different siRNAs. Y-axis is the ratios of RNAs RPM values (reads per million) in siRNA KD vs. siRNA control, e.g.: ($RPM_{siFBL-1}$+50) / ($RPM_{siCtrl}$+50). Each value was added 50 to avoid the division-by-zero problem and reduce variation. Note, this result is based on RPM, reads per million, so the actual magnitude of difference should be multiplied by the fold changes observed in Fig. 4F. For example, a 2-fold difference plotted here means that the fragment is up by 2*X fold, where X is the fold change in total tRF levels observed on the gel in Fig. 4.
(K-L) Scatter plots of tRNA fragment (tRF) levels between siCtrl and siFBL. Shorter fragments (less than 60% of full lengths) are plotted in red dots, while longer fragments are plotted in blue. The 2-dimensional error bars indicate standard deviations in siCtrl and siFBL, respectively. Shorter tRFs are labeled by name and start-end positions (e.g., Met_CAT (33,75), for 3' end tRNA half), for those that differ by >=0.05 between the two samples, or if their shorter fragment fractions are larger than 0.15. Linear regression statistics for the shorter and longer fragments are listed in the plot. Panel K shows the full distribution, while panel L shows a zoom-in view for tRFs with relative tRF abundance between 0 and 0.2.

**Figure S6. Characterization of D97/D133 KO HEK293 cells**
(A-B) Pearson correlation between HEK293 WT and D97/D133 single and double KO RNA-seq (A) and Ribo-seq (B). Replicates were merged before the scatterplot. r is Pearson correlation coefficient. n is the number of genes/RNAs detected at >=10 reads.
(C-D) Pearson correlation for 48 nuclear-encoded tRNA types between HEK293 D97/D133 single and double KO clones in total RNA-seq (C) and Ribo-seq (D). r is Pearson correlation coefficient. n is the number of genes/RNAs detected at >=10 reads.
(E) Redrawn plots from Bornelov et al. Fig. 2C,F [24], to highlight the amino acid usage similarities and differences with human stem cell differentiation studies. In particular, Met was not consistently down in the two experiments in both self-renewal and differentiation, different from our observation in HEK293 cells. Lys and Ile are consistently down, like our HEK293 D97/D133 KO cell lines. Pro is consistently up also like the D97/D133 KO cell lines. This comparison confirms that Met amino acid usage downregulation is unique to the D97/D133 double KO, and not a secondary effect of cell state transition.
(F) tRNA anticodons associated with differentiation and proliferation states, from Gingold et al. 2014 Table S2, and colored based on A/U or G/C content at the wobble position [25]. Note, the wobble position is the first nucleotide in the tRNA anticodon, but the third in the codon. The p-value was calculated with a chi-square test of enriched tRNAs between differentiation and proliferation states, where the null hypothesis is G/C and A/U starting anticodons should be equally enriched between the two cellular states.
(G) For nuclear-encoded mRNAs in RNA-seq and ribo-seq from WT and D97 and D133 single KO HEK293 cells, ratios of expression levels were ranked. Then the usage of all 66 codons (standard 64 plus initiator Met and selenocysteine) were calculated for the top 10% (most up-regulated) and bottom 10% (most down-regulated) mRNAs and weighted by expression levels. The ratios were plotted in ranked order. Blue: A/U-ending codons: Red: G/C ending codons. Asterisk (*): 3 stop codons. M and m: elongator and initiator methionine. U_uga: selenocysteine. This analysis shows that the elongator Met-AUG codon is only reduced on the translatome level, but not the transcriptome level.
(H) Same as panel G. except that amino acid usage was calculated for RNA-seq and ribo-seq data.
(I) Changes in codon usage frequencies were calculated and ratios KO/WT were plotted for RNA-seq and ribo-seq. The linear regression properties are at the bottom: equation, correlation coefficient r and statistical significance p. NN[GC]: G/C ending codons. NN[AU]: A/U ending codons. The inset ratios indicate ratios of average NN[GC] vs. average NN[AU] in RNA-seq and ribo-seq data. p values after the ratios are two-sided unpaired t-tests between the two codon groups.
(J) The following changes in amino acid (aa) usage frequencies were calculated between D97/D133 single KO HEK293 lines and wildtype (WT). aa freq. from RNA-seq: aa usage was calculated for 23 aa types, including the 20 standard amino acids plus initiator Met (m), selenocysteine (U) and stop (*), in mRNAs from KO and WT and the ratios were calculated and normalized so that median=1. aa freq. from Ribo-seq: same as the RNA-seq analysis, except that ribo-seq was used for the calculation. Standard deviations (s.d.) were calculated for the x and y-axes, horizontal and vertical, respectively. The linear regression properties are at the bottom: fit equation, correlation coefficient r and statistical significance p.

(K) Up and down regulated mRNAs based on log-transformed ratios between D97/D133 single KO vs. wildtype (WT) cells were median-normalized and tested for gene set enrichment using GSEA (c5.all collection, default parameters). Then enriched gene sets were clustered based on term similarity and tested for enrichment of GO terms using simplifyEnrichment. Only biological process terms (GOBP) were included in clustering, to avoid redundancy among the three major GO types: BP, CC (cellular components), and MF (molecular functions). Example GOBP term clusters that are consistent across RNA-seq, ribo-seq and GSCU analysis were highlighted in blue boxes and linked with gray lines. Similar enriched GO terms between the D97 and D133 KO cell lines were linked by gray curves.
(L) Same as panel K, except that the analysis was performed on ribo-seq mRNA levels.
(M) Same as panel K, except that the analysis was performed on translation efficiency (TE, i.e., mRNA levels in ribo-seq normalized by mRNA levels in RNA-seq).
(N-O) Example gene ontology (GO) terms overrepresented in high and low Met_AUG codons, from the analysis of APPRIS collection of principal transcripts, among GO terms in c5.all.v2023.1.Hs.symbols.gmt. BP: biological processes. CC: cellular components. (N)ES: (normalized) enrichment score. nom p: nominal p values.
(P) For each mRNA, the ratio of RNA-seq and ribo-seq levels in KO vs. WT HEK293 cells was calculated, and plotted in log10 scale. Then numbers of mRNAs distributed to the 4 quadrants were counted for selected GO terms. mito-mRNA: mRNAs encoded by the mitochondrial genome (12 detected out of 13 total). Oxphos: HALLMARK oxidative phosphorylation (GO:M5936, 57 detected out of 200 total). HALLMARK glycolysis (GO:M5937, 30 detected out of 200 total). Note, the standard ribosome profiling protocol used in this study is not very accurate for monitoring translation of mitochondria-encoded mRNAs. The HPG-labeling experiment in Fig. 5 provides an alternative confirmation for reduced translation of mitochondria-encoded proteins.
(Q) Gene specific codon usage (GSCU) values for Met AUG codon in the 5 complexes of oxidative phosphorylation, only showing nuclear-encoded mRNAs. Principal isoforms of human mRNAs in the APPRIS collection were used for calculation. Numbers in parentheses are the genes included in each complex. Not all components are enriched in Met.

**Figure S7. Characterization of D133 KO mESC**.
(A) Genomic DNA PCR validation of the three clones of D133 KO mES cell lines.
(B) Bright field view of WT and D133 KO TC1 mES and EBs at 4x and 20x magnifications.
(C) Reproducibility of RNA-seq from mES, EB and CM stages of WT and D133 KO cells. Raw reads were plotted in equal aspect of x-y axes. Pearson's correlation coefficient r and numbers of RNAs measured with >=10 reads are listed.
(D) Expression dynamics of snoRNAs and nuclear-encoded cytoplasmic tRNAs are measured in the RNA-seq data and shown in violin plus box plots. For each RNA, the WT mES value was set to 1. P values are from Wilcoxon signed rank test.
(E) Analysis of tRNA levels between D133 KO clone #5 and WT mES cells at different stages of differentiation. All nuclear-encoded tRNA genes were grouped by anticodons and then log2 transformed ratios between KO and WT were normalized so that median=1 for each KO/WT comparison and then plotted. The primary mapped location on a tRNA gene was counted for all multi-mapped reads.
(F) Up and down regulated mRNAs based on log-transformed ratios between EB and mES stages during stem cell differentiation for the WT cells were median-normalized and tested for gene set enrichment using GSEA (m5.all collection, default parameters). Then enriched gene sets were clustered based on term similarity and tested for enrichment of GO terms using simplifyEnrichment. Only biological process terms (GOBP) were included in clustering, to avoid redundancy among the three major GO types: BP, CC (cellular components), and MF (molecular functions).
(G) Same as panel F, except that the analysis was performed on CM vs. mES stage.
(H) Example GO BP term upregulated in WT CM vs. ES stage
(I) Expression differences between D133 KO and WT, and dynamics across ES, EB, and CM, for one-carbon metabolism mRNAs. Violin and box plots of the log transformed KO/WT ratios are summarized in the inset panel, showing the increased one-carbon metabolism activity in mES cells after D133 KO, which then returned to the same level as WT at the CM stage.
(J) The one-carbon cycles redrawn from Ducker and Rabinowitz et al. 2017, where upregulated enzymes are highlighted in red [26].

**Supplementary Tables**

**Table S1. All human snoRNA-target interactions captured by PARIS2.**
Fields include duplex group (DG), reads_number, chromosome (chr)_snoRNA, start_snoRNA, end_snoRNA, strand_snoRNA, chr_target, start_target, end_target, strand_target, Nm_pos, base-pairing, predicted secondary structure model, minimal free energy (MFE, kcal/mol), Box motif. DG information: RNA1,RNA2,DGID,covfrac, where covfrac, coverage fraction, is defined as connections between the two RNAs, divided by the square root of the product of all reads coverages at the the interacting regions (see details in [27]). n=7531 records, with at least one read support. Among all records, 6182 has no stable predicted structures (MFE=NA), 564 are with D box guides while 785 are with D' box guides. No H/ACA box elements were annotated due to difficulties in computational prediction. Interactions with annotated repetitive RNA genes beyond tRNAs include RN7SK, 47; RN7SL, 16; RNU7, 3; RNY, 8; U17, 12; hs45S, 3717; hs5S, 24; hssnRNA (U1, U2, U4, U5, U6, U11, U12, U4atac and U6atac), 347.

**Table S2. Comparison of PARIS2 data with known human rRNA Nm sites from Erales et al. 2017 and Yi et al. 2021.**
The GSE105248_known_Nm (Erales et al. 2017) data sheet contains 98 known sites from snoRNABase, separated into 3 types: guide snoRNA known in snoRNAbase (n=89), guide snoRNA unknown in snoRNAbase (n=6), and known Nm sites without PARIS support (n=3). The GSE159004_known_Nm (Yi et al. 2021) data sheet is the same except that the 5.8S sites were not measured. The GSE105248_with_sig_pvalue data sheet contains. The 45S pre-rRNA unit contains 18S (range 3655-5523), 5.8S (6601-6757) and 28S (7925-12994) subunits, based on GenBank entry NR046235.

**Table S3. Nm site analysis from HeLa siFBL, PCa siEZH2, HEK293 siFBL and snoKO cell lines.**
RMscore analysis based on data from HeLa siFBL (Erales et al. 2017) [2], PCa siEZH2 (Yi et al. 2021) [4], and HEK293 siFBL and D97/D133 double KO cell lines (this study). Previous publications used the original RMS method, whereas the new data in this study were produced using the dRMS method. n=40701 nucleotide positions were analyzed in the following RNAs: 5.8S, 18S, 28S, U1, U11, U12, U2, U3, U4, U4atac, U5, U6, U6atac, and 430 nuclear-encoded tRNA genes. Samples include: HeLa siCtrl and siFBL,

PCa C4-2 siCtrl and siEZH2, HEK293 WT, siCtrl, siFBL, D97/D133 double KO. Additional fields include, chromosome-position (chr_pos), RNA name, position on RNA, adjusted positions for the standardized 73nt tRNA when applicable (aligned_Pos), Nm levels in siCtrl (WT) and siFBL samples, potential guide snoRNAs, minimal free energy (MFE). In HEK293 dRMS data, only the 3' end reads were used for the analysis.

**Table S4. Human PARIS2 and PARCLIP gapped reads supporting snoRNA-tRNA interactions.**
Total reads for PARIS2: 1550. Total reads for PAR-CLIP: 630. n=954 for all interactions with at least 1 read support from either method. For the tRNA multi-gene families, each family is represented once. Members of snoRNA families are presented individually due to higher sequence variation (e.g., the SNORD116 family).

**Table S5. Predicted and PARIS2/PARCLIP-derived snoRNA-tRNA interactions.**
All predicted human snoRNA-tRNA interactions, n=44540, and the following details: snoRNA_name, tRNA_name including genomic location in hg38, full-length snoRNA_seq, full-length tRNA_seq, gudie_box, guide_seq, tRNA_snoRNA_duplex, tRNA_snoRNA_structure, Nm_pos, predicted minimal free energy (mini_MFE), MFE after shifting the target site to the left and right in single nt steps for 30 nucleotides (shift_MFE), PARIS_reads and CLIP_reads supporting the interactions. For multi-gene families, each snoRNA and each tRNA is represented in separate records.

**Table S6. Nm site analysis for yeast ncRNAs using published RMS data.**
The following genes are analyzed: 18S, 25S, 5.8S, 5S, and 267 tRNA genes, n=26748 records. Fields include: WT replicates 1-3 and bcd1 D72A mutant replicates 1-3 from Khoshnevis et al. 2021., WT replicates 1-3 and Dbp7KO replicates 1-3 from Aquino et al. 2021 Nature Comm., WT replicates 1-2 and Dbp3KO replicates 1-2 from Aquino et al. 2021 NAR. In particular, tRNA positions were lifted to standard nomenclature (length=73). For each sample, 5end_coverage, 3end_coverage, total_coverage and Nm levels were calculated (wherever applicable).
